# Proteomic signature of the Dravet syndrome in the genetic *Scn1a*-A1783V mouse model

**DOI:** 10.1101/2021.04.27.441099

**Authors:** Nina Miljanovic, Stefanie M. Hauck, R. Maarten van Dijk, Valentina Di Liberto, Ali Rezaei, Heidrun Potschka

**Author notes:** Correspondence:* Dr. H. Potschka, Institute of Pharmacology, Toxicology, and Pharmacy, Ludwig-Maximilians-University, Koeniginstr. 16, D-80539 Munich, Germany. Declarations of interest: none.

## Abstract

**Background:** Dravet syndrome is a rare, severe pediatric epileptic encephalopathy associated with intellectual and motor disabilities. Proteomic profiling in a mouse model of Dravet syndrome can provide information about the molecular consequences of the genetic deficiency and about pathophysiological mechanisms developing during the disease course.

**Methods:** A knock-in mouse model of Dravet syndrome with *Scn1a* haploinsufficiency was used for whole proteome, seizure and behavioral analysis. Hippocampal tissue was dissected from two-(prior to epilepsy manifestation) and four-(following epilepsy manifestation) week-old male mice and analyzed using LC-MS/MS with label-free quantification. Proteomic data sets were subjected to bioinformatic analysis including pathway enrichment analysis. The differential expression of selected proteins was confirmed by immunohistochemical staining.

**Results:** The findings confirmed an increased susceptibility to hyperthermia-associated seizures, the development of spontaneous seizures, and behavioral alterations in the novel *Scn1a*-A1873V mouse model of Dravet syndrome. As expected, proteomic analysis demonstrated more pronounced alterations following epilepsy manifestation. In particular, proteins involved in neurotransmitter dynamics, receptor and ion channel function, synaptic plasticity, astrogliosis, neoangiogenesis, and nitric oxide signaling showed a pronounced regulation in Dravet mice. Pathway enrichment analysis identified several significantly regulated pathways at the later time point, with pathways linked to synaptic transmission and glutamatergic signaling dominating the list.

**Conclusion:** In conclusion, the whole proteome analysis in a mouse model of Dravet syndrome demonstrated complex molecular alterations in the hippocampus. Some of these alterations may have an impact on excitability or may serve a compensatory function, which, however, needs to be further confirmed by future investigations. The proteomic data indicate that, due to the molecular consequences of the genetic deficiency, the pathophysiological mechanisms may become more complex during the course of the disease. Resultantly, the management of Dravet syndrome may need to consider further molecular and cellular alterations. Ensuing functional follow-up studies, this data set may provide valuable guidance for the future development of novel therapeutic approaches.

## Introduction

In the vast majority of patients with Dravet syndrome, an *SCN1A* mutation can be identified, which results in functional deficiency of the encoded sodium channel subunit Nav1.1 (Brunklaus and Zuberi, 2014; Dravet and Oguni, 2013). Dravet syndrome is characterized by seizures with a poor pharmacoresponse to available antiepileptic drugs (Wallace et al., 2016). Moreover, there is a high risk of sudden unexpected death in epilepsy (SUDEP) in Dravet patients (Kalume, 2013). Thus, despite the licensing of orphan drugs for Dravet syndrome, there is an unmet clinical need for novel therapeutic approaches tailored to the disease. While knowledge about the genetic cause provides a basis for the rational development of precision medicine approaches (Dugger et al., 2018), the elucidation of the molecular consequences of the genetic deficiency can further improve our understanding of the pathophysiological mechanisms involved, thus providing a broader framework for the identification of novel targets and the subsequent development of innovative therapeutic strategies.

Large-scale proteomic profiling constitutes one of the most promising tools providing comprehensive information about epilepsy-associated alterations at a functionally relevant molecular level. During recent years, initial studies have identified proteome alterations in models of acquired epilepsy (Bitsika et al., 2016; Keck et al., 2017; Keck et al., 2018; Li et al., 2010; Liu et al., 2008; Walker et al., 2016). However, respective data for genetic epilepsies are rather limited; they focus on models of genetic absence epilepsy and fragile X syndrome (Danış et al., 2011; Liao et al., 2008; Xu et al., 2018). To our knowledge, the molecular consequences of *Scn1a* genetic deficiency are yet to be studied by quantitative whole proteome analysis. Characterizing the molecular signature of Dravet syndrome may therefore contribute to the understanding of disease-associated alterations in neuronal homeostasis.

Here, we completed a large-scale proteomic profiling study in a novel conditional mouse line carrying a human Dravet syndrome *SCN1A* mutation (Kuo et al., 2019; Ricobaraza et al., 2019). The analysis focused on two time points: prior to and following the onset of spontaneous recurrent seizures. This design obtained information about the molecular patterns occurring both during epileptogenesis and following epilepsy manifestation.

Taken together, our findings improve the understanding of the complex molecular consequences of *SCN1A* genetic deficiency and provide a basis for the discovery of novel innovative targets, both for the prevention of disease progression and therapeutic management of Dravet syndrome.

## Material and methods

### Genetic mouse model: breeding and genotyping

Breeding colonies of the parental lines B6(Cg)-*Scn1a*^*tm1*.*1Dsf*^/J (#026133(Kuo et al., 2019; Ricobaraza et al., 2019)) and 129S1/Sv-*Hprt*^*tm1(CAG-cre)Mnn*^/J (#004302(Tang et al., 2002)) were generated based on breeding pairs purchased from the Jackson Laboratory (Bar Harbor, Maine, USA). Conditional knock-in male mice with a floxed *Scn1a* (mutation A1783V in exon 26) were crossed with female mice heterozygous for Cre recombinase (X-linked to neuronal promoter *Hprt* gene). The offspring were heterozygous Dravet mice carrying the A1783V mutation (wildtype or heterozygous for Cre recombinase) or wildtype mice without the A1783V mutation (wildtype or heterozygous for Cre recombinase). Since the presence of Cre did not affect the phenotype, animals were divided into a Dravet group and a wildtype control group depending on the presence of the A1783V-*Scn1a* mutation. This mouse line was chosen for the experiments because A1783V represents one of the clinically relevant mutations affecting S6 transmembrane region of domain IV of the alpha subunit of type I voltage-gated sodium channels in patients with Dravet syndrome (Lossin, 2009). The genotype of the animals was confirmed by PCR, using 5′-GCAACTCTTCACATGGTACTTTCA-3′, 5’-GCACCTCTCCTCCTTAGAACA-3’ and 5′-GGAGAAACACGAGCAGGAAG-3′ primers: wildtype, 164 bp; heterozygous pre Cre, 164, 198 and 410 bp; and heterozygous post Cre, 164 and 198 bp (Fig. 1A). The presence of Cre recombinase in mice was also confirmed by PCR, using 5′-CTGGTGCTTTACGGTATCGC-3′, 5’-TTCATAGAGACAAGGAATGTGTCC-3’ and 5′-AATCCAGCAGGTCAGCAAAG-3′ primers: WT allele, 217 bp; Cre allele, 450 bp.

**Fig. 1.**
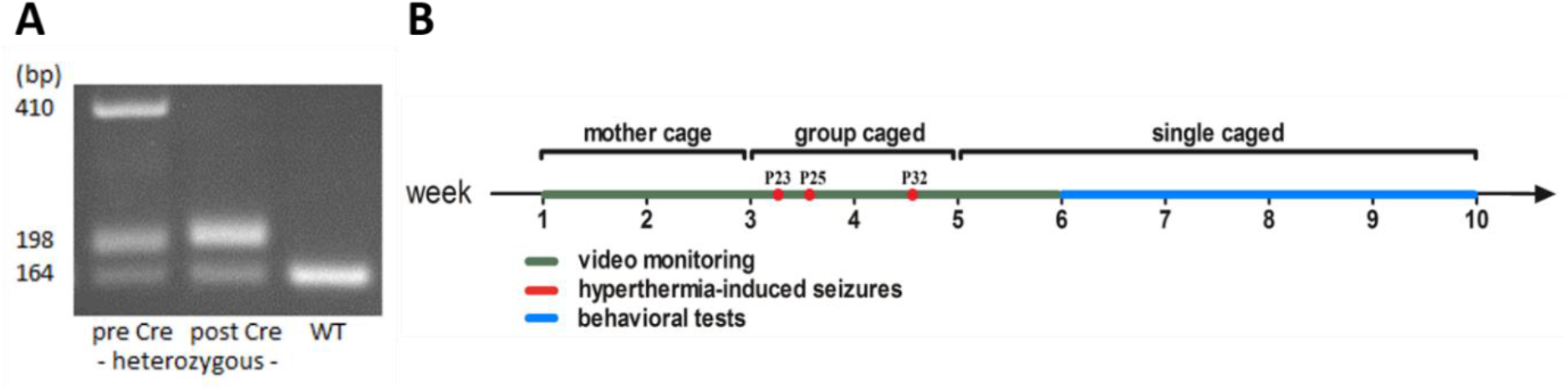
**A** PCR genotyping for distinguishing Dravet mice with a Cre activated A1783V mutation (post Cre heterozygous) and wildtype mice. **B** Experimental timeline.

Experiments were approved by the responsible government of Upper Bavaria (license number 55.2-1-54-2532-166-2015 and 55.2-1-54-2532-168-2016). All experiments were conducted in accordance with the EU directive 2010/63/EU for animal experiments and the German Animal Welfare act. All experiments were planned and carried out considering the ARRIVE guidelines and the Basel declaration (http://www.basel.declaration.org) including the 3R concept.

The brain tissue from twenty mice was used for proteomic analysis with five animals per genotype and time point. Sections from four wildtype and four Dravet mice aged 4 weeks were also used for the qualitative immunohistochemical analysis of selected proteins. An additional group of animals was used for the phenotypic characterization of Dravet mice. From 38 heterozygous mutant mice, 23 (11 male; 12 female) survived the phase around weaning, which was characterized by a high seizure-associated mortality rate. Data from these animals were compared with those from 21 wildtype littermate controls (11 male; 10 female). One wildtype and one Dravet female mouse from this group were later used for EEG-telemetry recordings.

In addition, baseline data from 22 heterozygous mutant mice were considered for seizure monitoring based on parallel video monitoring and telemetric EEG recordings. These animals were also prepared for ECG recordings and were further used in a separate study assessing the effects of ketogenic diet in Dravet mice (Miljanovic et al., under revision).

### Housing of animals

Each litter was housed in individually ventilated cages (Tecniplast, Hohenpeißenberg, Germany) from birth until weaning (3 weeks). Following this phase, mice were housed 3-5 animals per standard Makrolon type III cage (Ehret, Emmendingen, Germany). In order to confirm the presence of seizures, animals used for brain tissue sampling and proteomic analysis were single housed in Makrolon type II open cages following weaning (Ehret, Emmendingen, Germany). Animals used for behavioral assessments were single housed 2 weeks prior to behavioral assessment. The order of cages in the rack was randomized (randomizer.org).

Each cage was supplied with two nestlets (Ancare, Bellmore, New York, USA), one animal house (Tecniplast, Hohenpeißenberg, Germany; Zoonlab GmbH, Castrop-Rauxel, Germany) and fresh sawdust as bedding material once per week (Lignocel, Rosenberg, Germany). Standard conditions in the animal facility were: lights on from 6 am to 6 pm, temperature 22 ± 2° C, and humidity 50 ± 10 %. Animals received food (ssniff® R/M-H, Sniff, Soest, Germany) and water *ad libitum* with Dietgel76A offered as a supplement (Sniff, Soest, Germany) between postnatal days P14 and P26.

### Brain samples for proteomic analysis

Brain samples were obtained at two different time points from five mice per genotype and time point. Two-week-old mice (early time point, prior to seizure onset) were sacrificed by decapitation whilst the four-week-old mice (later time point) were sacrificed by cervical dislocation. Hippocampal tissue from the left hemisphere was dissected and fresh frozen in liquid nitrogen (Besamungsstation München-Grub, Poing, Germany) using 1.5 ml Protein LoBind Tubes (Eppendorf, Wesseling-Berzdorf, Germany. The experimenter was blinded to animal genotype and the order of animals for dissection was randomized (randomizer.org). Samples were frozen and stored at -80 °C until analysis. The right brain hemisphere was collected and left in 4% PFA solution for 24 hours and then switched to 30 % sucrose solution, all at 4 °C. The brain was cut into 40 µm coronal sections, which were later used for immunohistochemistry.

### Proteome analysis

Untargeted proteome analysis was carried out by an experimenter not involved in the *in vivo* experiments to avoid any expectation-triggered bias. Snap frozen hippocampus samples were directly bead milled with a Precellys homogenizer (Peqlab, Lutterworth, U.K.) in extraction buffer (10 mM Tris-HCl pH 7.6 with 1 % NP40, 10 mM NaCl and complete protease inhibitors) as previously described (Molin et al., 2015). The total protein concentration in the sample was measured by the Bradford assay. A modified filter-aided sample preparation (FASP) method was used for the digestion of 10 µg of total protein per sample as described (Lepper et al., 2018). A Q Exactive HF mass spectrometer was used for proteomic screening (ThermoFisher Scientific, Dreieich, Germany) with operation in the data independent acquisition (DIA) mode (Lepper et al., 2018). Per sample, one injection unit of the HRM Calibration Kit (Biognosys, Schlieren, Switzerland, #Ki-3003) was used for spiking 1 µg of peptides for indexing of retention time. Samples were loaded automatically onto the UPLC system (Ultimate 3000, Dionex, Sunnyvale, CA). The system contained a nano trap column (inner ∅ = 300 μm × 5 mm, packed with Acclaim PepMap100 C18, 5 μm, 100 Å; LC Packings, Sunnyvale, CA). Following 5 minutes of elution from the trap column, the peptides were separated by reversed-phase chromatography (Acquity UPLC M-Class HSS T3 Column, 1.8 μm, 75 μm × 250 mm; Waters, Milford, MA) using a 7−27 % gradient of acetonitrile (flow rate of 250 nL/ minute, 90 minutes) followed by two short gradients of 27−41 % acetonitrile (15 minutes) and 41−85 % acetonitrile (5 minutes). After 5 minutes at 85 % acetonitrile, the gradient was reduced to 3 % acetonitrile over 2 minutes and then allowed to equilibrate for 8 minutes. All acetonitrile solutions contained 0.1 % formic acid.

The DIA method comprised alternating mass spectrometry (MS) full scans spanning from 300−1650 m/z at 120,000 resolution, followed by 37 DIA window scans at 30,000 resolution for peptide fragmentation with a variable width ranging from 300−1650 m/z. Normalized collision energy was adjusted to 28, with profile type spectra recording.

The DIA liquid chromatography with tandem MS (LC−MS/MS) raw files were converted (HTRMS converter) and analyzed (Spectronaut version 11, Biognosys, Schlieren, Switzerland) as described (Lepper et al., 2018). An automatic calibration mode was chosen with precision indexed retention time (iRT) alignment enabled for the application of the nonlinear iRT calibration strategy. Peptides were identified by comparison with an in-house accumulated spectral library, which has been obtained from mouse brain samples measured on the same MS set-up with a data-dependent acquisition mode. Peptide identification was filtered for a false discovery rate (FDR) of 1 %. Only proteotypic peptides were considered for quantification of proteins, applying MS2 area based summed precursor quantities. The data filtering function was set to q-value percentile mode applying a 50 % setting, thus enabling a match between runs. This setting allows only peptide precursor signals passing the FDR threshold of 1 % in over 50 % of all samples to be further considered for identification and quantification, thus working towards a reduction of false positive identifications. The sum of abundances of all unique peptides per protein was log2 transformed and obtained values were compared between groups using an unpaired Student’s t-test with a significance level of p<0.05.

### Pathway enrichment analysis

Pathway enrichment analysis was completed using a publicly available pathway tool (Consensus PathDB over-representation tool, (Kamburov et al., 2009)). A background list was used, comprising of all identified proteins. Only pathways with both a p<0.01 and at least two dysregulated proteins were considered. Only pathways reaching a q<0.01 (=p-value corrected for multiple testing using the FDR method) were considered significant and discussed in this study. Protein abundances of all proteins assigned to significantly changed pathways were visualized in a heat map. Individual fold changes for each animal were calculated by dividing their value against the wildtype group’s mean and log2 transforming this value. The resulting matrix was visualized using R software version 3.5.1 (Team, 2017) and “gplots” package (Warnes et al., 2016).

### Immunohistochemical staining of PPP1R1B (DARPP-32)

In order to further confirm expression alterations of selected proteins, brains from four-week-old wildtype and Dravet mice (n=4 per group) were used for immunohistochemistry. Free-floating sections were washed in PBST (phosphate-buffered saline, 0.1% Tween 20) at room temperature and heat-induced epitope retrieval (HIER) was performed at 80 °C for 30 minutes using sodium citrate buffer (pH 6.0). Sections were cooled down on ice and rinsed in PBST. Endogenous peroxidase was inhibited (3 % H_2_O_2_ in TBS for 60 minutes). Slices were rinsed in PBST and a blocking step in 6 % goat serum (60 minutes) was performed to prevent non-specific antibody binding. Sections were then incubated overnight at 4 °C with a monoclonal rabbit anti-DARPP32 primary antibody (Abcam, Berlin, Germany, Cat# ab40801, lot# GR3213231-3) diluted 1:2500. The next day, sections were washed in PBST and incubated for 60 minutes at room temperature with a secondary biotinylated goat anti-rabbit antibody (Vector laboratories, Cat# BA-1000, lot# 2F0430) diluted 1:1000. After washing steps in TBST, brain sections were incubated at room temperature in VECTASTAIN ABC-Peroxidase Kit (Vector Laboratories Cat# PK-4000, RRID: AB_2336818, lot#2337238) for 60 minutes, diluted 1:100. The slices were washed in PBS and stained using SIGMAFAST™ 3,3′-Diaminobenzidine tablets (Sigma-Aldrich, Darmstadt, Germany, Cat# D4418, lot# SLBR2966V) for 1 minute.

Brain sections were quickly rinsed in distilled H2O, washed in PBS, mounted on microscope glasses using PBST, and cover slipped with Entellan® (107960, Merck, Darmstadt, Germany). Negative controls were processed in parallel without the primary antibody.

### Immunohistochemical staining of HSD11B1

The staining protocol for HSD11B1 was identical to the one described above (DARPP-32) with the following exceptions. Inhibition of endogenous peroxidase was done in 3 % H_2_O_2_ in TBS for 15 minutes. A blocking step was performed with 1.5 % goat serum for 120 minutes. The primary antibody was anti-HSD11B1 (Abcam, Berlin, Germany; Cat# ab39364, lot# GR3247054-7) diluted 1:200. Slices were stained in SIGMAFAST™ 3,3′-Diaminobenzidine solution for 100 s.

### Microscopy

Bright field images were captured at 4x, 10x and 40x magnification with an Olympus BH2 microscope with a single chip charge-coupled device (CCD) color camera (Axiocam; Zeiss, Göttingen, Germany) and an AMD AthlonTM 64 processor based computer with an image capture interface card (Axiocam MR Interface Rev.A; Zeiss, Göttingen, Germany).

### Hyperthermia-induced seizures and threshold determination

Hyperthermia-induced seizures were analyzed in mice at postnatal day 23, 25, and 32. Mice were transported to the laboratory 30 minutes prior to seizure induction. Temperature and light intensity in the laboratory were adjusted to 22 ± 2° C and 600 lux. On all experimental days, tests began at 12 p.m. and the order of animals was randomized (randomizer.org). Observers were blinded to animal genotype as far as possible. At the early time point, blinding was difficult considering the obvious phenotype (lower body weight). The whole procedure was video recorded (Axis communications, Lund, Sweden). Body temperature was measured continuously with a RET-4 rectal probe (Physitemp, Clifton, New Jersey, USA), which was placed in warm saline solution (B. Braun Vet Care GmbH, Tuttlingern, Germany) before use. Subjects were placed in a plexiglass cylinder for 5 minutes to habituate to the environment and to record basal body temperature data. The IR lamp connected to the temperature controller (Physitemp, Clifton, New Jersey, USA) was then turned on to slowly increase the body temperature with a ramping rate of 0.5° C per 2 minutes (Oakley et al., 2009). Heating was stopped immediately when generalized tonic-clonic seizures were observed or when the body temperature reached 42°C. Animals were allowed to cool down before returning them to their home cage. All equipment was cleaned with 70 % ethanol between subjects (CLN, Langenbach, Germany).

The severity of hyperthermia-induced seizures was assessed based on the Racine scoring system as follows: I (mouth and facial movements), II (head nodding), III (forelimb clonus), IV (rearing with forelimb clonic convulsions) and score V (generalized clonic convulsions followed by rearing and falling) (Racine, 1972).

### Spontaneous seizures

Information about spontaneous seizure onset, frequency, duration, and severity score was obtained via continuous video monitoring from the second postnatal week for 6 weeks. Videos were carefully reviewed by experienced technicians at a maximum of 8x playback speed and all generalized seizures were documented. Two eight-month-old female mice (one wildtype, one Dravet) and 22 twelve-week-old Dravet mice (11 males, 11 females) underwent survival surgery for telemetry (ETA-F10 or HD-X02, DSI, St. Paul, USA) and EEG electrode implantation. The order of animals was randomized (randomizer.org). Thirty minutes before anesthesia induction, mice received 1 mg/kg meloxicam s.c. (Metacam®, Boehringer Ingelheim, Germany). For general anesthesia, mice received 400 mg/kg chloral hydrate i.p. (Carl Roth, Karlsruhe, Germany)(n=2) or isoflurane (Isofluran CP®, Henry Schein Vet, Hamburg, Germany) with a concentration of 4 % and 1.5 % for anesthesia induction and maintenance, respectively (n=22). The local anesthetic bupivacaine (0.5 %; Jenapharm®, Mibe GmbH, Brehna, Germany) was applied subcutaneously to surgical areas affected by transmitter implants and leads placement. For intracranial electrode placement, bupivacaine with epinephrine (0.5 % + 0.0005 %; Jenapharm®, Mibe GmbH, Brehna, Germany) was applied subcutaneously.

Firstly, the skin was opened in the dorsocaudal part of the scapula region for placing the telemetric transmitter subcutaneously. Mice were then fixed in a stereotactic frame and three screws were inserted into the skull. The negative EEG lead was connected to the screw over the cerebellum. The positive EEG lead was connected to a bipolar Teflon-isolated stainless-steel electrode, which was implanted into the CA1 region of the hippocampus (ap: -2,00; lat: + 1,3; dv: -1,6). The electrode was fixed with Paladur (Heraeus®, Hanau, Germany). The skin over the skull was closed with absorbable sutures, while the initial cut for the transmitter placement was closed with tissue adhesive (Surgibond®, Henry Schein Vet, Hamburg, Germany).

As stated above, 22 animals were prepared for additional ECG analysis in the context of another study. Therefore, the negative ECG lead was fixed intramuscularly to the right pectoral muscle, whilst the positive ECG lead was fixed left of the xyphoid process prior to electrode implantation. Skin over the ECG connections was closed with absorbable sutures (Smi AG, St. Vith, Belgium). Animals were given oxygen until regaining consciousness. On the following day, mice received 1 mg/kg meloxicam s.c. Mice were allowed to recover for a full 2 weeks followed by a two-week continuous recording phase. In parallel, animals were video monitored (Axis communications, Lund, Sweden) in order to confirm and analyze behavioral seizure activity. Data were acquired with Ponemah software (Ponemah R, v. 5.2.0, DSI, St. Paul, USA) and seizure activity was detected automatically (Neuroscore™ v. 3.0, DSI, St. Paul, USA).

### Behavioral characterization

Following hyperthermia-induced seizures, all animals were tested in different behavioral paradigms except for one female Dravet mouse and one female wildtype mouse because of an age difference compared with the remaining animals. Thus, 22 Dravet mice (11 males; 11 females) and 20 wildtype (11 males; 9 females) mice were used for behavioral assessments.

One male Dravet mouse died following a spontaneous seizure and was therefore not exposed to the elevated plus maze and accelerated rotarod test. Data from one male Dravet mouse were not considered in the saccharin preference test due to leakage of one of the water bottles. During all behavioral paradigms, animals were single-housed as a presupposition for the social interaction test. Social interaction was analyzed at an age of 7 weeks and the order of the subsequent tests was as follows: open field test, saccharin preference test, elevated plus maze and accelerated rotarod test (Fig. 1B). The testing was completed until an age of 10 weeks. The order of animals for each test was randomized (randomizer.org).

Nest-building activity as well as saccharin preference were assessed in the home cage. All other behavioral tests (social interaction, open field test, elevated plus maze, accelerated rotarod test) were completed in a test room (temperature 22 ± 2° C, humidity 55 ± 5 %) under different light conditions adjusted to the specific paradigm (stated below). All tests were carried out during morning hours (starting from 8 a.m.). All behavioral test runs were documented by photographing (nest complexity) or video-recording (all other tests).

### Open field test

The open field paradigm is a widely used to assess exploratory behavior and locomotion in an unfamiliar environment (Carola et al., 2002). Mice were placed in the test room for 1 hour prior to testing to habituate (lighting 15-20 lux). Two round shaped arenas (Ø 61 cm, height 40 cm) were simultaneously used for the test. Time spent in three different zones (wall, middle and center) was analyzed with Ethovision 8.5 software (EthoVision XT, RRID:SCR_000441). Each mouse was placed in one arena 10 cm away from and facing the wall. Locomotion was recorded for 30 minutes and rearing behavior was counted manually. After completing the test, mice were returned to their home cage. The arena was cleaned with 0.1 % acetic acid between animals.

### Saccharin preference test

The saccharin preference test was used to test anhedonia-associated behavior as one of the symptoms of depressive disorders (Klein et al., 2015). The analysis was conducted during four consecutive 24-hour-long periods in the home cage. Animals were provided with two water bottles. First, both bottles were filled with water to determine water intake over 24 hours. The bottle on the left side was then filled with a 0.1 % saccharin solution. During the next day, both bottles were again filled with water. On the last day, the right bottle was filled with 0.1 % saccharin solution in order to test for a potential side preference bias. Liquid consumption was measured after each period.

### Nest-building activity

During postnatal week five, mice were individually placed into type III open cages. The following week, animals were provided with two new nestlets and nest complexity was assessed on a daily basis. Nests were photographed each morning. The images were later scored by an investigator blinded to group allocation. Nest complexity was ranked according to the scoring system developed by Jirkof and colleagues as follows (Jirkof et al., 2013): score 0 = nestlet intact, possibly carried around the cage; score 1 = nestlet is poorly manipulated with more than 80 % of the nestlet intact; score 3 = evident nest site with most of shreds in the nest site, less than 80 % nestlet material intact, hollow in bedding and mice begin to build walls; score 4 = flat nest, hollow in bedding, walls are higher than mice and encasing the nest less than 50%; score 5 = complex, bowl-shaped nest with walls higher than mice and encasing the nest more than 50%.

### Social interaction test

The social interaction test was performed to assess autism-associated behavioral patterns and affinity towards interaction with another mouse. Mice were single-housed in type III open cages for 2 weeks prior to testing. On the first 2 days, animals were transported to the test room (lighting 15-20 lux) and placed in an empty type III open cage for 10 minutes to habituate. On the test day, animals were transported to the test room 30 minutes prior to testing. Two mice of the same sex, same genotype and approximately same weight were then simultaneously placed in an empty cage and left for 10 minutes. Active and passive social interaction were measured with a stopwatch by an observer blinded to animal genotype and sex. The cage was cleaned with 0.1 % acetic acid between subjects. Sniffing, grooming or following the partner as well as aggressive behavior were considered as active social interaction. Laying or sitting next to each other without any interaction was classified as passive social interaction (Holter et al., 2015).

### Elevated plus maze

The elevated plus maze is generally used to assess anxiety-like behavior in rodents (Ben-Hamo et al., 2016). Mice were placed in the test room for 1 hour prior to the experiment. The maze comprised of two open arms (40 cm long and 10 cm wide) and two closed arms (same dimensions with walls 15 cm high). The plus maze was elevated 68 cm above the floor. Light intensity was set to approximately 200 lux in the open and 60 lux in the closed arms. Animals were placed in the center of the maze facing the open arm. A five-minute-long trial was recorded with Ethovision XT for each animal. Head-dipping and stretching behavior (exploring the open arms with the head while the body stays in the closed arms or center part of the maze) were recorded manually by observers blinded to animal group allocation. The maze was cleaned with 0.1 % acetic acid between animals.

### Accelerated rotarod test

The accelerated rotarod test is widely used to evaluate motor performance in mice (Shiotsuki et al., 2010). Mice were placed in the test room for 1 hour prior to assessment to habituate (15-20 lux). A rotarod apparatus (Ugo Basile 47600, Varese, Italy) was used to assess the animals’ motor coordination. The rotation accelerated from 4 to 40 rpm over 5 minutes. Each animal was subjected to four consecutive trials with approximately 2 minutes break for cleaning the rod (0.1 % acetic acid). Testing was carried out on three subsequent days, the first for training and the others for testing. Male animals were always tested before females. Passive rotations, time remaining on rod and current rod speed were recorded.

### Statistical analysis

Statistical analysis was performed with R version 3.5.1. GraphPad Prism (Version 5.04 and 6.01, GraphPad, USA) was used for data visualization, except for heat maps, which were visualized with R, and Venn diagrams, visualized with an online tool (http://bioinformatics.psb.ugent.be/webtools/Venn/). The functional and molecular annotation of differentially expressed proteins was prepared using a Panther classification system (http://pantherdb.org/). Figure 8 was prepared using a pathway downloaded from KEGG PATHWAY Database (https://www.kegg.jp/kegg/pathway.html). A Spearman correlation matrix was calculated using R version 3.5.2. The significance level for correlation analysis was set at < -0.5 or > 0.5. Animals with missing data were not considered for the respective correlation analysis.

All data were expressed as mean ± SEM with the exception of nest complexity data, for which the median is illustrated in the respective graph. All results were first checked for possible batch effects. If present, batch effects were considered in the statistical analysis.

Two-tailed unpaired t-tests were used for comparison between experimental and control groups where indicated. Two-, three-, four- and five-way ANOVAs were used to test the effects of genotype, sex, time, batch and their interaction where appropriate. Comparisons, which included multiple measurements, were tested using repeated measures ANOVA. ANOVA tests were followed by a Bonferroni post-hoc test. A two-tailed Mann-Whitney non-parametric test was used for analyzing nest complexity scores. The significance level was set at p < 0.05 for all tests.

### Data availability

The raw data sheet with individual protein abundances, which supports the findings of this study, is provided in *Appendix B*. A complete list of abbreviations of significantly regulated proteins mentioned in the manuscript sorted by their function is provided in *Supplementary Material* (Table A.1).

## Results

### Breeding pattern and outcome

Twenty-three female mice heterozygous for Cre recombinase (129S1/Sv-*Hprt*^*tm1(CAG-cre)Mnn*^/J) were bred with 21 conditional knock-in male mice with floxed *Scn1a-1783V* (B6(Cg)-*Scn1a*^*tm1*.*1Dsf*^/J) in pairwise and trio matings split into two different batches. Eighteen of them delivered litters with a total of 38 heterozygous mutant mice (Dravet mice) and 41 wildtype littermates. The sex ratio in the offspring was almost balanced with 38 males (19 Dravet; 19 WT) and 41 females (18 Dravet; 23 WT). Due to the 40 % mortality rate around the time of weaning, 15 Dravet mice were lost. The remaining 23 Dravet mice and randomly selected 21 wildtype mice (randomizer.org) were used for model characterization.

### General condition, body weight development, seizure thresholds and spontaneous seizures

The body weight of Dravet mice proved to be significantly lower at the time point of weaning. However, following weaning, affected animals showed a good development of body conditioning scores finally reaching a comparable body weight to wildtype mice (Fig. 2A).

**Fig. 2.**
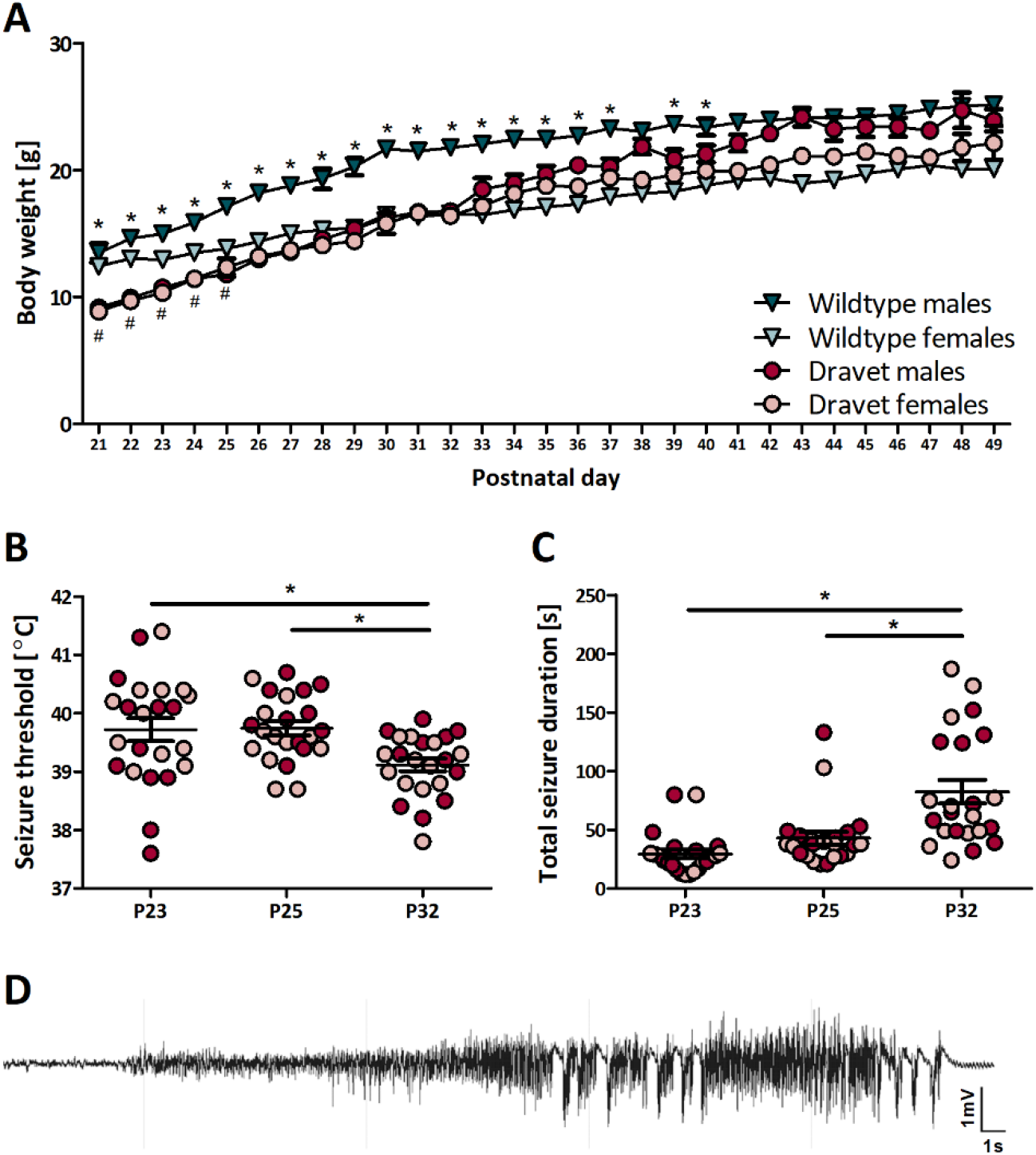
Body weight, spontaneous and hyperthermia-induced seizures. **A** Body weight development following weaning. Body weight in animals with a Dravet genotype (males n = 11; females n = 11) was significantly lower in the early phase following weaning (females until P25, males until P41) as compared to wildtype mice (males n = 11, females n = 9) (Unpaired t-test; * p < 0.05 males, # p < 0.05 females, mean ± SEM). **B** Seizure threshold temperature at P23, P25, and P32. The threshold significantly decreased on P32 compared to previous testing. **C** Total duration of hyperthermia-induced seizure activity, calculated as the sum of all motor seizure periods occurring immediately following stimulation. The duration significantly increased with repeated stimulations. **B-C** Data are from 11 male and 12 female animals with a Dravet genotype (Two-way RM ANOVA, Bonferroni post hoc; * p < 0.05, mean ± SEM**). D** Representative EEG recording of a generalized tonic-clonic seizure (Racine score V, followed by running und bouncing) in an adult female mouse.

In response to hyperthermia induction, all Dravet mice exhibited generalized tonic-clonic seizures on all three testing days. In contrast, their wildtype littermates did not exhibit motor seizure activity despite increasing their body temperature to at least 41 °C.

The number of Dravet mice showing running and bouncing behavior increased with repeated hyperthermia induction reaching 18/23 animals on P32. The average threshold temperature for seizure induction was 39.7 ± 0.9 °C at P23, 39.7 ± 0.6 °C at P25, and 39.1 ± 0.5 °C at P32 (mean ± SD, Fig. 2B). Seizure duration significantly increased with subsequent stimulations (Fig. 2C). No differences were observed when comparing female and male mice.

Video monitoring of experimental animals in their home cages demonstrated the development of spontaneous motor seizures at P16. The seizures observed included generalized tonic-clonic seizures, sometimes associated with running and bouncing indicating seizure spread towards the brain stem. In addition, prolonged phases with behavioral arrest, immobility, and lack of responsiveness to external stimuli were observed. Between P20 and P23 several Dravet mice died. In several instances, video monitoring indicated that the death was directly associated with a generalized seizure in these animals thus indicating probable SUDEP. The mortality rate reached 40 %. The remaining 23 Dravet animals were used for behavioral characterization. SUDEP or probable SUDEP was rarely observed in older animals. Only one animal was found dead in its cage following the fourth postnatal week.

To further confirm spontaneous seizure activity, telemetric EEG recordings were performed with simultaneous video recordings in an adult female Dravet and female wildtype mouse. During the one-week recording, the Dravet mouse exhibited multiple generalized tonic-clonic seizures often followed by running and bouncing. Assessment of the EEG recordings confirmed electrographic seizure activity with high amplitude spiking over 500 µV (Fig. 2D). In the wildtype mouse, no evidence was obtained for electrographic seizure events.

Additional recordings in a group of three-month old Dravet mice confirmed a high penetrance of the epilepsy phenotype with multiple generalized tonic-clonic seizures in 19/22 mice. On average, mice experienced seven seizures per week (range: 2 to 16; data not shown). Seizures frequently occurred in clusters with animals often exhibiting seizures on only two subsequent days per week (range: one to four, data not shown). The average seizure duration was 50.19 s (data not shown).

### Phenotype: behavioral alterations

In the open field paradigm, hyperlocomotion was evident in male and female Dravet mice. Throughout the 30-minute test, the total distance moved and the rearing frequency was significantly greater in Dravet mice as compared to wildtype animals (Fig. 3A-B). Moreover, immobility time was proved to be shorter in Dravet mice (data not shown). Additionally, thigmotaxis proved to be increased in Dravet mice with more time spent in the wall zone. Time spent in the other zones (middle and center) was not affected by the genotype (Fig. A.1C).

**Fig. 3.**
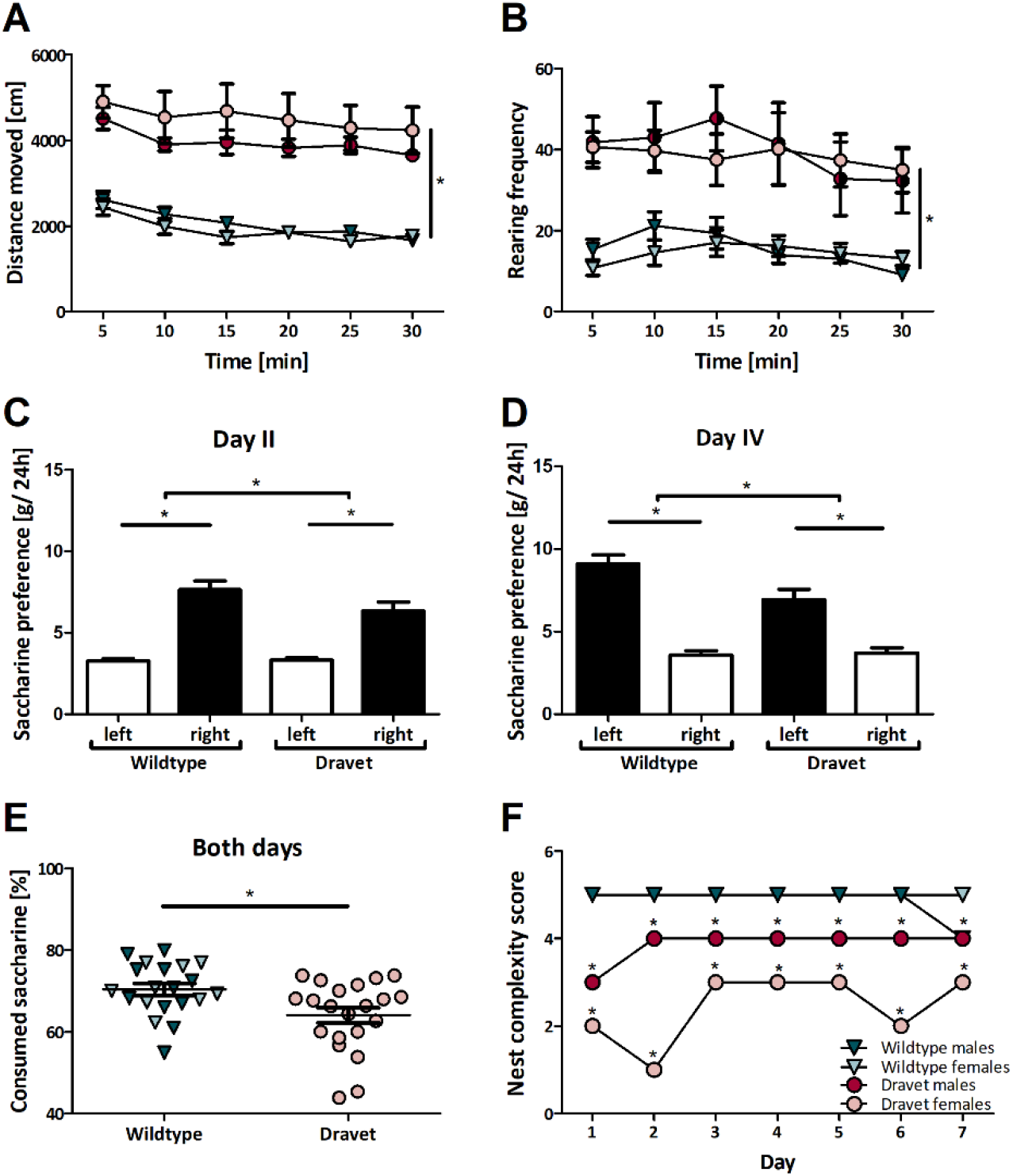
Open field test, saccharin preference and nest-building behavior. **A** Distance moved in the open field test over 30 minutes, divided into mean intervals of 5 minutes. The total distance moved of Dravet mice significantly exceeded that of wildtype mice. **B** Rearing frequency. Dravet mice exhibited more frequent rearing positions than wildtype mice. **A-B** Data are from 22 animals with a Dravet genotype and 20 wildtype animals (Three-way RM ANOVA, Bonferroni post hoc; * p < 0.05, mean ± SEM). **C-D** Water or saccharin solution consumption per 24 h. Both Dravet and wildtype mice preferred saccharin solution over water, independent of a side preference. **E** Total saccharin solution consumption over both days. Dravet mice consumed significantly less saccharin solution as compared to wildtype mice. **C-E** Data shown are from 21 animals with a Dravet genotype and 20 wildtype animals (Three-way ANOVA, Bonferroni post hoc; * p < 0.05, mean ± SEM). **F** Nest complexity score over 7 days. As shown in the graph, nests of Dravet mice (n=22) received lower scores as compared to those from wildtype mice (n=20)(Mann-Whitney non-parametric test; * p < 0.05, median).

Regardless of genotype, a preference for saccharin solution over water was observed. However, the amount of saccharin consumed by wildtype mice exceeded that of mice with the Dravet genotype. In line with this finding, the percentage of consumed saccharin solution was reduced in Dravet mice (mean ± SD: Dravet mice 64.07 ± 8.64 %; wildtype mice 70.38 ± 6.41 %) (Fig. 3E).

Interestingly, when saccharin consumption was compared between the first and the second exposure, Dravet mice consumed similar amounts whereas wildtype mice consumed more of saccharin solution on the subsequent exposure (Fig. 3C-D). Findings were comparable in male and female mice.

Nest-building behavior, a non-essential activity, demonstrated a poorer performance in mice with a Dravet genotype (Fig. 3F).

When analyzing social interaction following a period of social isolation in single-housing, all Dravet mice spent more time engaged in active and less time in passive social interaction when compared to wildtype littermates (Fig. A.1A-B).

In the elevated plus maze, Dravet mice spent more time in aversive parts, i.e. the open arms of the maze, compared to the wildtype animals (Fig. A.1D). In addition, a higher frequency of head dips and a reduction in stretching behavior was evident in all Dravet mice (Fig. A.1E-F). Sex differences were not observed.

We applied the accelerated rotarod test to address disturbances in motor coordination. During habituation, animals from both groups showed a “learning” curve with an improvement in performance with subsequent trials. Thereby, it was evident this was related to an increased focus of the animals on the task. When compared to wildtype mice, Dravet mice remained on the rod for longer. Additionally, females performed better than males in both the control and experimental groups (Fig. A.1G-I).

The Spearman correlation coefficients between selected parameters were calculated. Open field test variables including total distance moved showed a significant correlation (all p<0.001) with social interaction (active r=0.69, passive r=-0.52), nest complexity score (r=-0.57), time spent in aversive parts (r=0.58), and number of head-dips in the elevated plus maze test (r=0.76).

### Proteomic profiling

Proteomic profiling identified over 4000 different proteins in the mouse hippocampus samples. Comparison of protein abundance revealed that, as a consequence of the genetic *Scn1a* deficiency, significant alterations in the expression of 205 and 881 proteins were observed in two- and four-week-old Dravet mice as compared to wildtype littermates, respectively (unpaired t-test, p<0.05, Fig. 4A). While the majority of these differentially expressed proteins were up-regulated at the early time point, more proteins were down-regulated at the later time point (Fig. 4B). As the down-regulation of proteins can be a general consequence of neuronal damage and cell loss, expression of the neuronal marker NeuN was assessed. It remained at control levels for both time points (data not shown). Moreover, we confirmed that the heterozygous loss-of-function *Scn1a* mutation did not result in changes in NaV1.1 protein abundance regardless of the time point (data not shown). Although changes in protein expression as a consequence of a missense mutation are not necessarily expected, this finding is of relevance for model characterization as both enhanced degradation of a non-functional protein as well as a compensatory expressional up-regulation are possible.

**Fig. 4.**
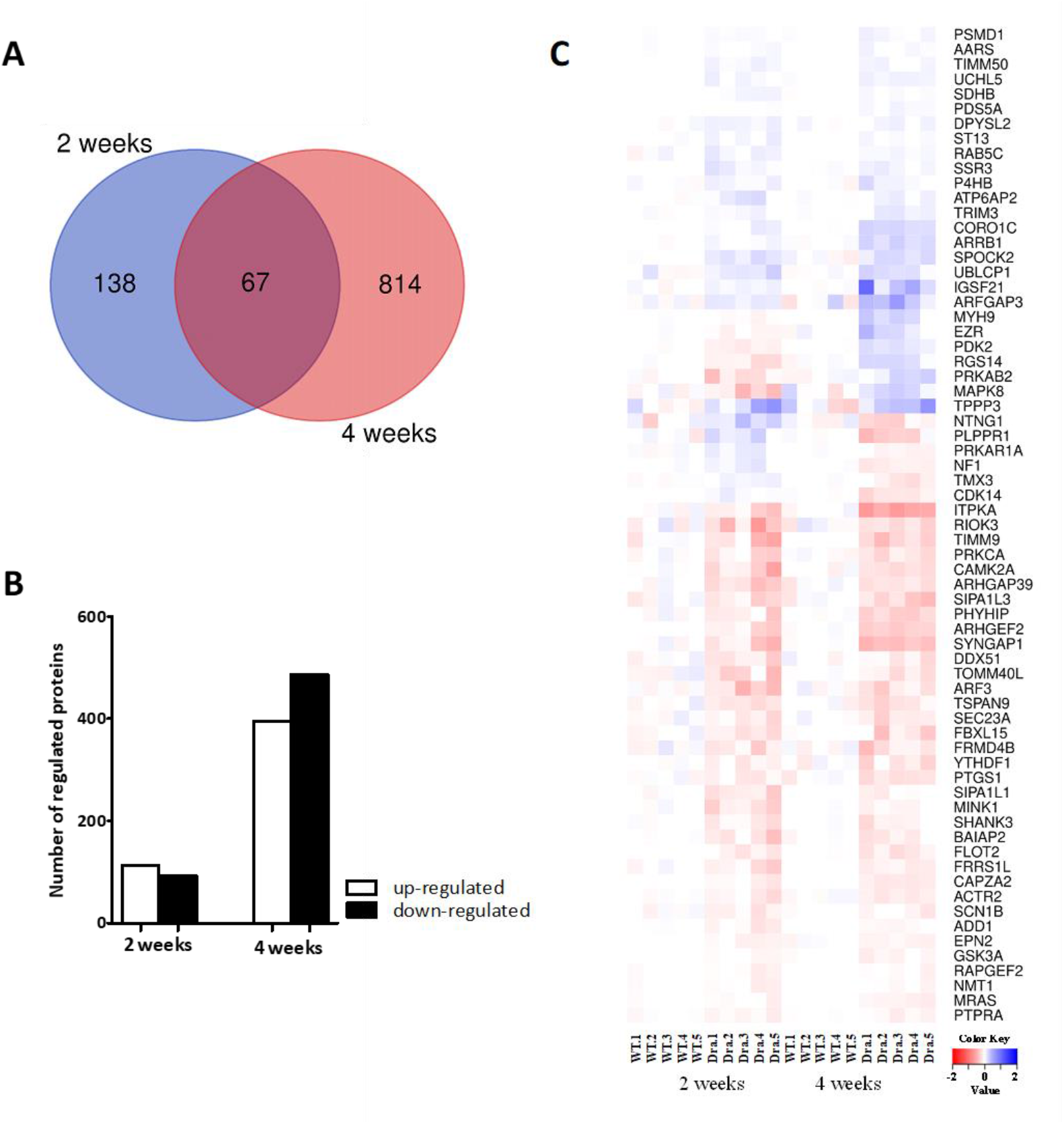
Differentially expressed proteins in Dravet mice. **A** Overlap of differentially expressed proteins in two- and four-week-old Dravet mice illustrated by a Venn diagram. **B** Total number of differentially expressed proteins in two- and four-week-old Dravet mice. **C** The heat map illustrates the fold change in protein level in relation to the mean of the wildtype group in two- and four-week-old Dravet mice. The respective color key is provided beneath the heat map.

A direct comparison of the datasets from both time points revealed an overlap of 67 differentially expressed proteins (Fig. 4A). Most of the proteins maintained the direction of change. However, some proteins were down-regulated at the earlier time point but later showed an overexpression or vice versa (Fig. 4C).

While a more pronounced proteome alteration was evident at the late time point, the functional annotation of differentially expressed proteins from both time points revealed some similarities in the qualitative pattern of protein regulation with a comparable distribution of regulated proteins to different functional groups (Fig. 5A).

**Fig. 5.**
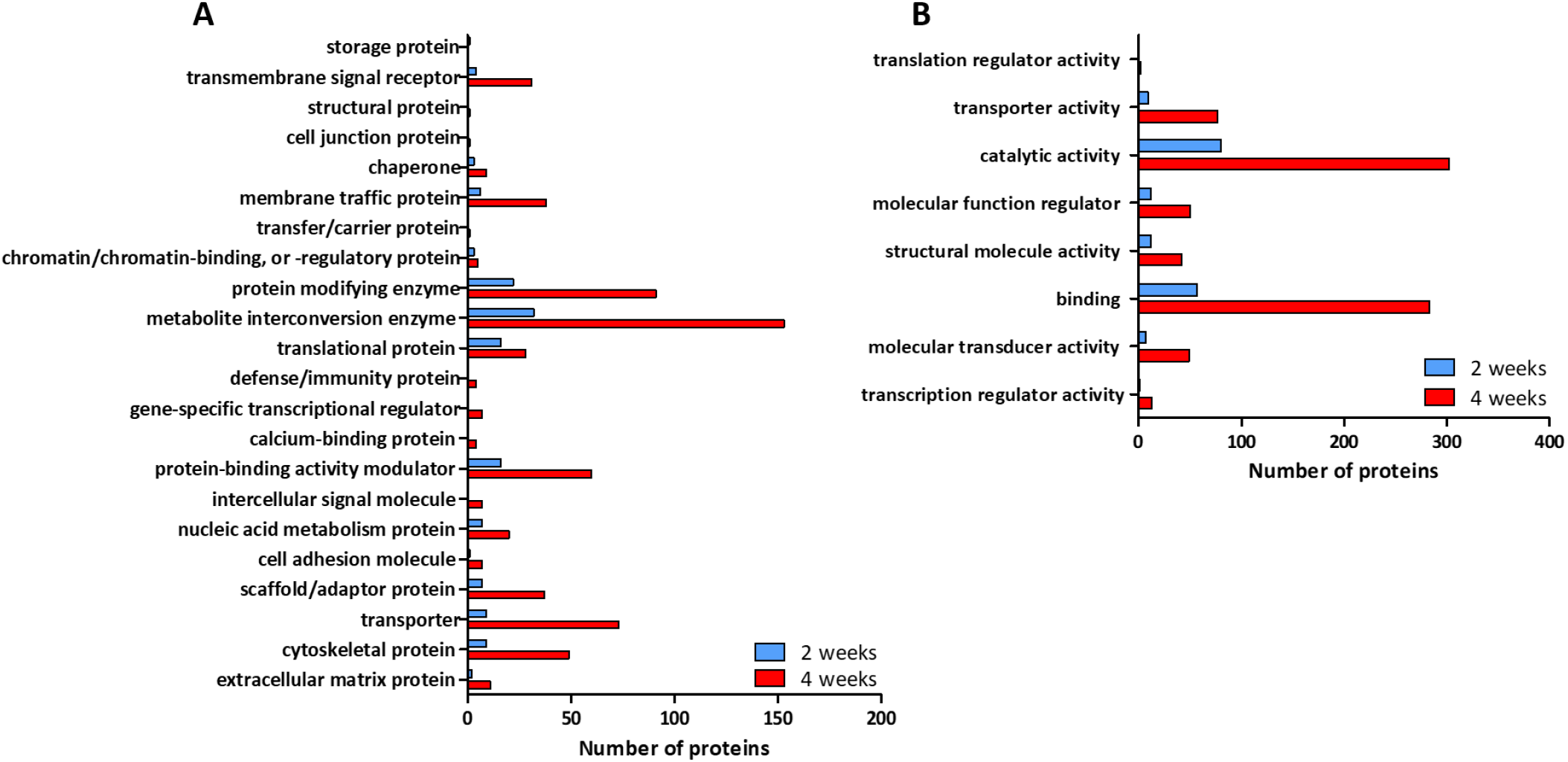
Classification of differently expressed proteins. Functional (**A**) and molecular (**B**) annotation of differentially expressed proteins in two- and four-week-old Dravet mice.

The majority of regulated proteins were classified as metabolite interconversion enzymes, protein modifying enzymes, protein-binding activity modulator and translational proteins at the earlier time point, and metabolite interconversion enzymes, protein modifying enzymes and transporters at the later time point. Regarding the molecular function of dysregulated proteins, again, a more pronounced protein regulation was evident at the later time point (Fig. 5B). Proteins associated with catalytic activity and binding exhibited the strongest regulation at both time points.

### Differential Protein Expression – early time point

Proteomic profiling in Dravet mice prior to the occurrence of first spontaneous seizures and epilepsy manifestation can provide information about the process of epileptogenesis as a direct consequence of *Scn1a* genetic deficiency. As mentioned above, a pathway enrichment analysis was completed to obtain general information about the regulation pattern concerning function and neurobiological significance. Pathway enrichment analysis did not identify any significantly regulated pathway (q<0.01). When considering significantly regulated individual proteins in Dravet mice (unpaired t-test, p<0.05), the most prominent down-regulation became evident for Ras-specific guanine nucleotide releasing factor 1 (RASGRF1). RASGRF1 is a member of the Ras GTP protein family, which plays a role in synaptic plasticity (Brambilla et al., 1997). RASGRF1 is associated with NMDA receptors (Krapivinsky et al., 2003) and its regulation or dysfunction has been discussed in the context of epileptogenesis and epilepsy manifestation (Chen et al., 2018; Tonini et al., 2001; Vlaskamp et al., 2019).

Considering the more modest regulation of most other significantly regulated proteins, we will highlight a selection based on their putative functional relevance and a minimum change of at least 15 % compared to expression rates in wildtype mice (Fig. 6A).

**Fig. 6.**
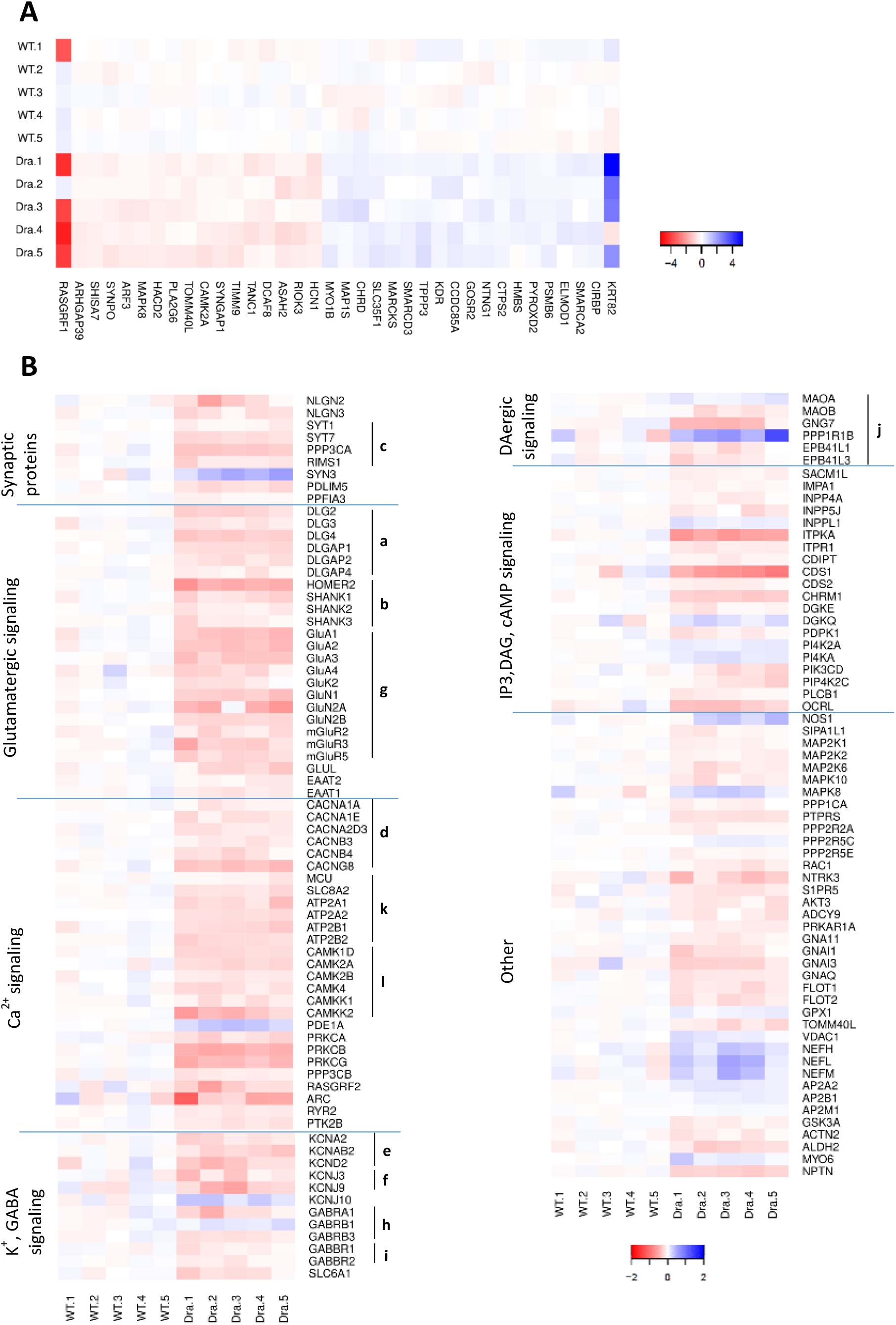
Expression analysis of proteins significantly regulated in Dravet mice before and following epilepsy manifestation. **A** Expression analysis of proteins significantly regulated in Dravet mice prior to epilepsy manifestation with a minimum change of at least 15 % compared to expression rates in wildtype mice. The R package ‘gplots’ was used to generate heat maps. The heat map illustrates the fold change in protein level in relation to the mean of the wildtype group and the respective color key is provided next to the heat map. **B** Expression analysis of proteins linked to all significantly enriched pathways in Dravet mice following epilepsy manifestation. The R package ‘gplots’ was used to generate heat maps. The heat map illustrates the fold change in protein level in relation to the mean of the wildtype group and the respective color key is given underneath the heat map. **a** PSD membrane-associated guanylate kinase protein family, **b** PSD scaffolding proteins, **c** proteins associated with synaptic vesicles, **d** voltage-gated calcium channel proteins, **e** voltage-gated potassium channel proteins, **f** inward-rectifier potassium channel proteins, **g** glutamatergic receptor proteins, **h** GABAA receptor subunits, **i** GABAB receptor subunits, **j** proteins functionally associated with dopaminergic (DAergic) synapse function, **k** calcium transporter proteins, **l** calcium/calmodulin-dependent protein kinases. (Blue cell color indicates an up-regulation, while red cell color stands for a down-regulation).

CAMK2A encodes the alpha subunit of calcium/calmodulin-dependent protein kinase and represents a key player in synaptic plasticity (Lisman et al., 2002). Two-week-old mice exhibited a reduced expression rate that persisted at the four-week time point.

Additionally, Dravet mice showed an up-regulation of VEGF receptor KDR (VEGFR2). It is linked with tight junction disassembly and blood-brain barrier dysfunction, which in turn can contribute to epileptogenesis (Morin-Brureau et al., 2012).

Cytidine triphosphate synthetase 2 (CTPS2) is a protein mediating CTP synthesis from UTP, a process that is linked with glutamine deamination to glutamate (Kassel et al., 2010). In two-week-old Dravet mice, we obtained evidence for an increased expression level of CTPS2.

### Differential Protein Expression – later time point

Proteomic profiling following epilepsy manifestation capturing direct and indirect consequences of *Scn1a* deficiency in Dravet mice revealed 881 regulated proteins (unpaired t-test, p<0.05) and 42 regulated pathways (q<0.01, Table 1). A heat map, which illustrates the level of individual protein change in relation to the mean of the wildtype group, is presented in Fig. 6B.

**Table 1.**
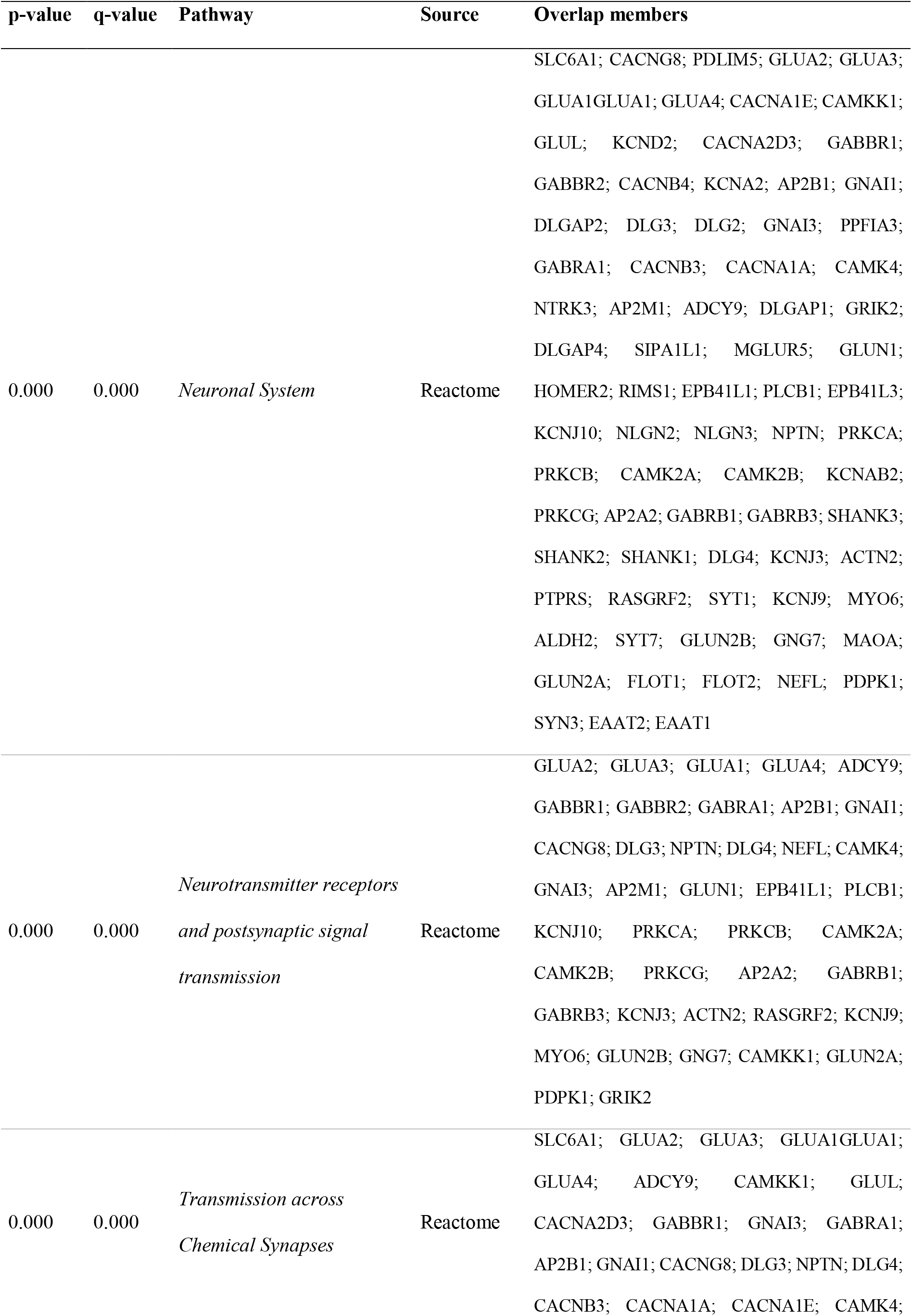
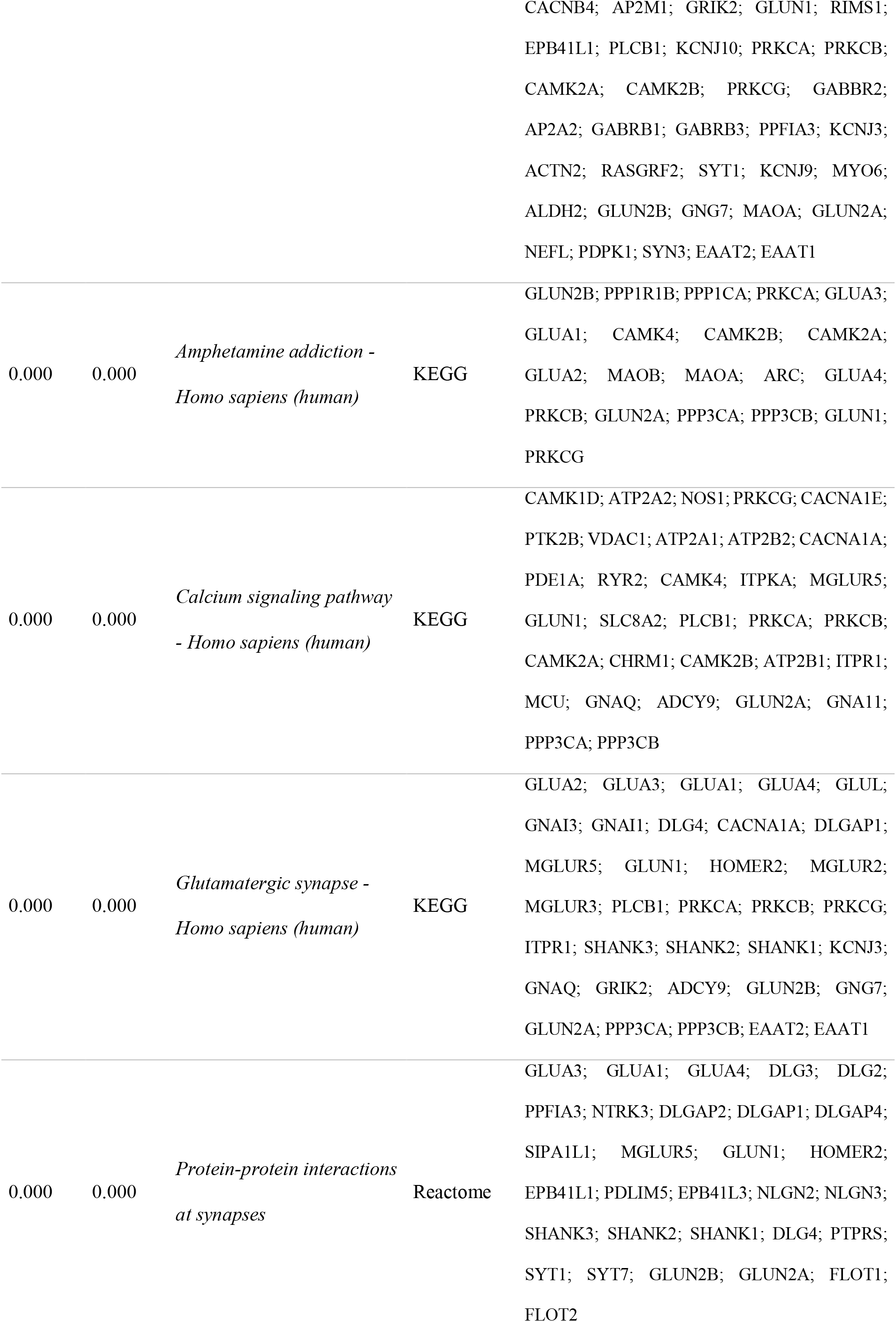
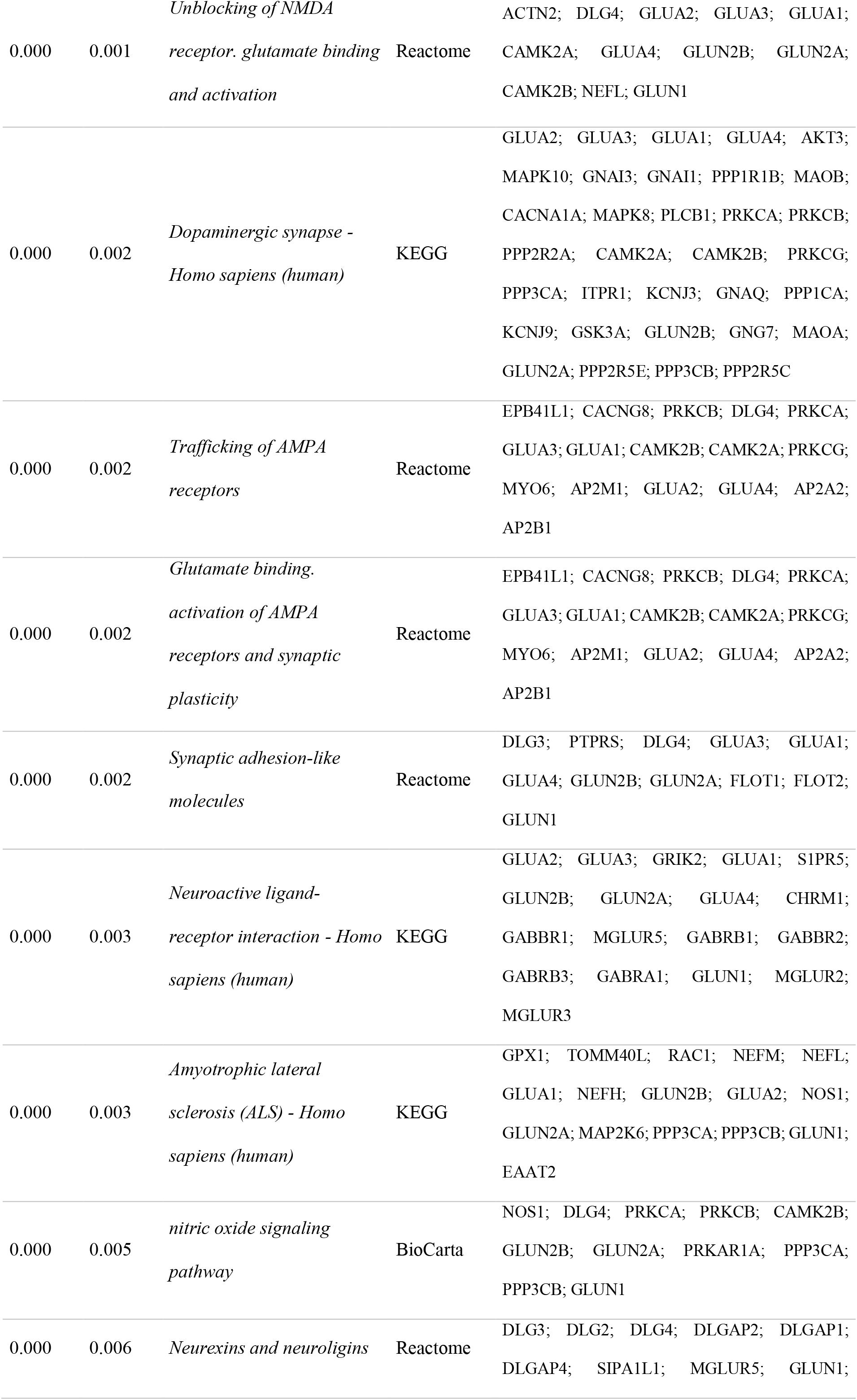
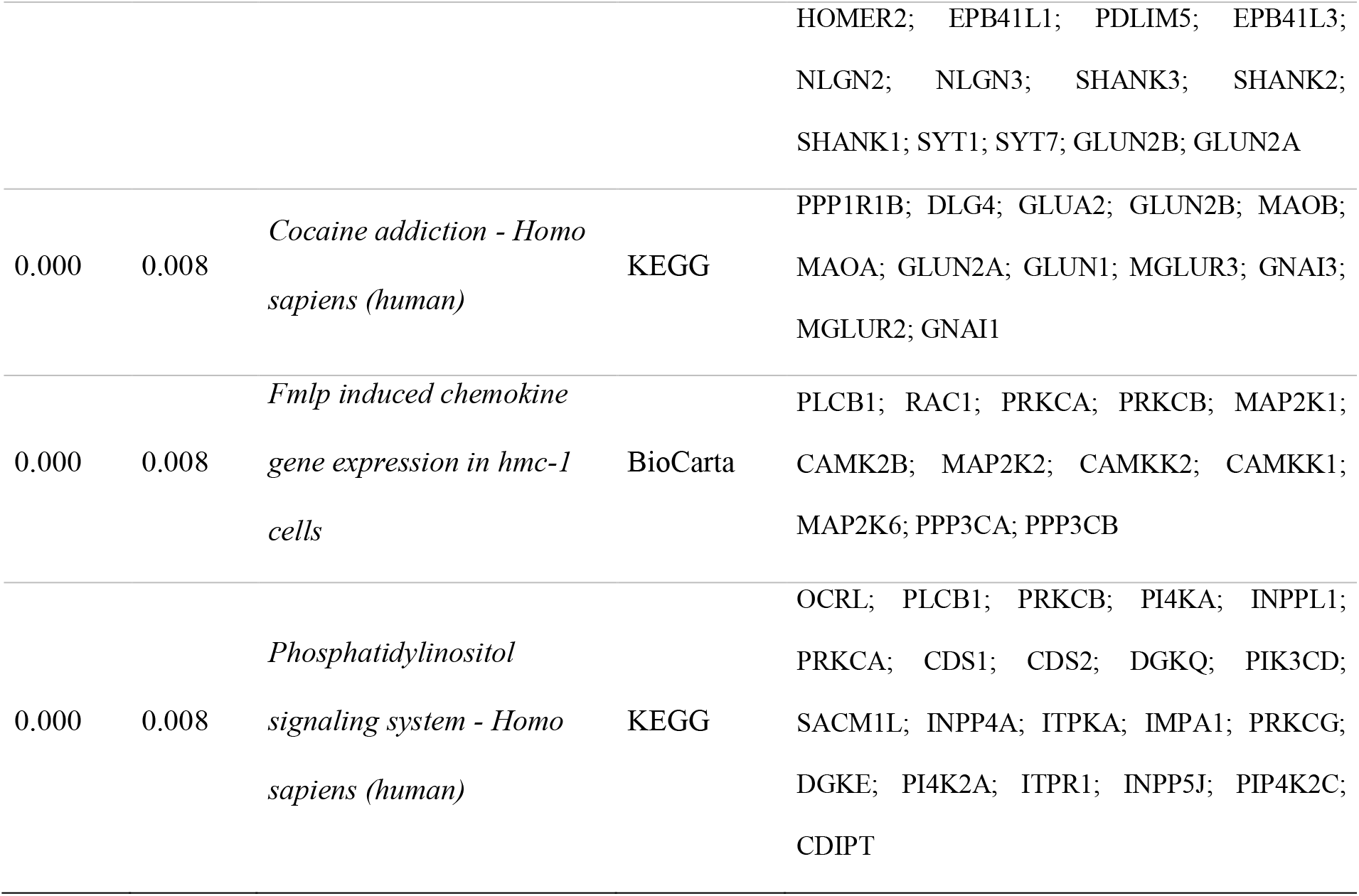
Overrepresented pathways in four-week old mice (q<0.01, ConsensusPathDB)

**Table 2.**
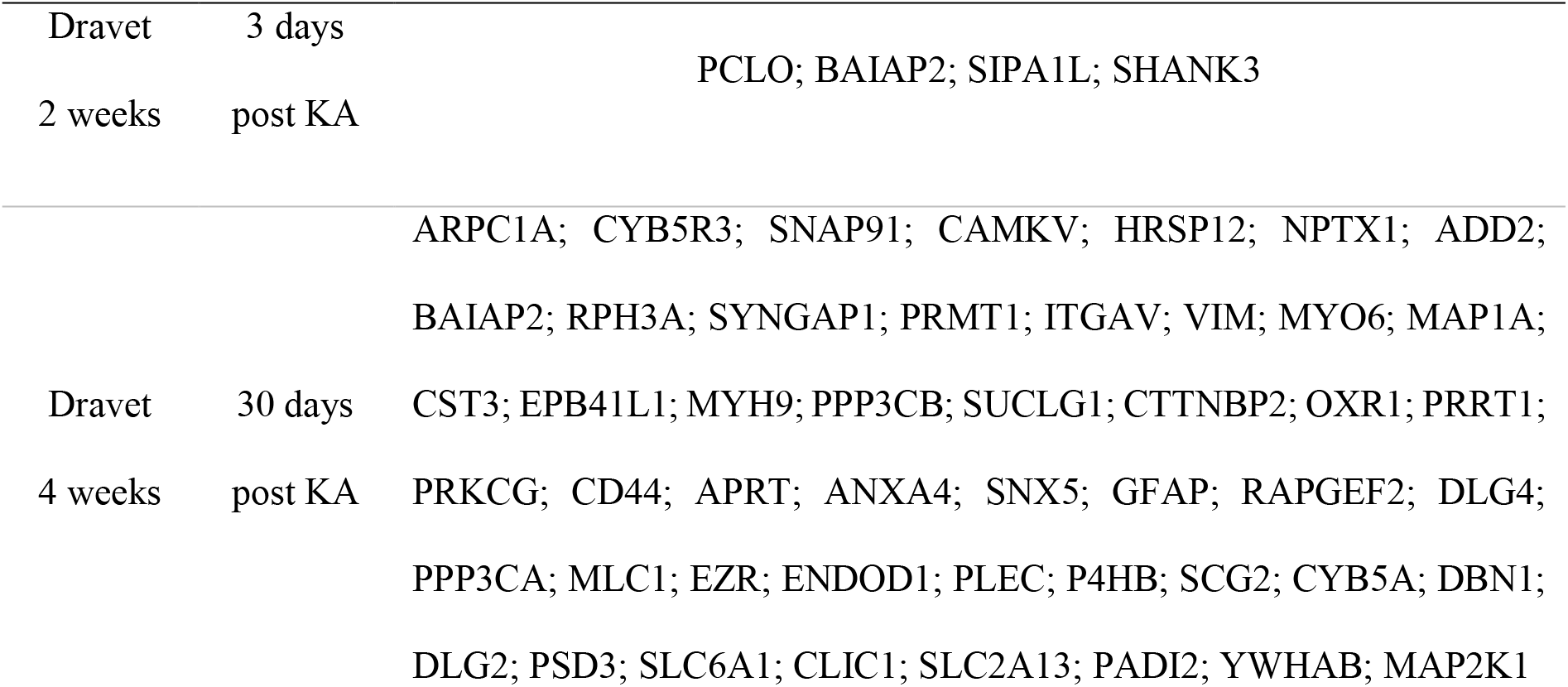
An overlap in differentially expressed proteins with a kainic acid mouse model of mesiotemporal lobe epilepsy reported by *Bitsika et al*., *2016*.

Pathways functionally linked to synaptic transmission dominated the list of significantly enriched pathways, in total comprising five pathways associated with general synaptic function and its regulation (*Neurotransmitter receptors and postsynaptic signal transmission*; *Transmission across Chemical Synapses*; *Protein-protein interactions at synapses*; *Synaptic adhesion-like molecules* and *Neurexins and neuroligins* with corresponding q-values 4.14e^-05^; 4.14e^-05^; 2.33e^-04^; 0.002; 0.006). The list of differentially expressed proteins linked with these pathways comprised 68 proteins. Several of these proteins proved to be down-regulated as a consequence of the genetic deficiency. These include synaptic adhesion-like molecules such as neurexins and neuroligins (Fig. 6B: NLGN2, NLGN3), which mediate trans-synaptic signaling and facilitate the processing of complex signals in neuronal networks (Südhof, 2008), as well as postsynaptic density (PSD) proteins including proteins of the membrane-associated guanylate kinase protein family (Fig. 6B-a), and scaffolding proteins (Fig. 6B-b). Moreover, these pathways included proteins associated with synaptic vesicles (Fig. 6B-c).

Proteins linked with ion channel function showed a complex regulation pattern in Dravet mice. The abundance of voltage-gated calcium channels (Fig. 6B-d), voltage-gated potassium channels (Fig. 6B-e) and two inward-rectifier potassium channels (Fig. 6B-f) was reduced in Dravet mice. Another inward-rectifier potassium channel (Fig. 6B: KCNJ10) was up-regulated in Dravet mice.

Interestingly, several neurotransmitter receptor proteins exhibited a differential expression pattern in Dravet mice. The list of these proteins was dominated by glutamatergic receptor proteins (Fig. 6B-g), which were all expressed at lower levels in Dravet mice. In addition, a change in expression of three GABAA receptor subunits (Fig. 6B-h) and of two GABAB receptor subunits (Fig. 6B-i) was evident with an induction of GABRA1 and GABBR1, and a down-regulation of GABRB1, GABRB3 and GABBR2.

In the context of neurotransmitter signaling, it is of additional interest that four pathways involved in glutamatergic signaling, specifically AMPA and NMDA receptor activation, binding, and synapses were regulated in Dravet mice (*Glutamatergic synapse*; *Unblocking of NMDA receptor, glutamate binding and activation*; *Trafficking of AMPA receptors*; *Glutamate binding, activation of AMPA receptors and synaptic plasticity* with q-values 1.4e^-04^; 0.001; 0.002; 0.002, respectively). Thereby, Dravet mice exhibited a reduced expression of iono- and metabotropic glutamate receptors, glutamate transporters (Fig. 6B: EAAT1, EAAT2), and glutamine synthetase (Fig 6B: GLUL).

Pathway enrichment analysis also revealed an overrepresentation of proteins functionally associated with dopaminergic synapse function (q=0.002, Fig. 6B-j). The changes in the pathway were dominated by alterations in the expression of proteins regulating dopamine metabolism or stabilizing D2 and D3 receptors on plasma membranes with MAOA up-regulated and MAOB, EPB41L1, EPB41L3 down-regulated. In four-week-old Dravet mice, reduced expression levels were also evident for GNG7, the G protein responsible for A2A adenosine or D1 dopamine receptor-induced neuroprotective responses (Schwindinger et al., 2012). Another protein that plays a role in synaptic plasticity and can be modulated by both dopaminergic D1 and glutamatergic NMDA receptors is the neuronal phosphoprotein PPP1R1B. Our data set revealed an overexpression in the hippocampus of four-week-old Dravet mice, which was further confirmed by immunohistochemistry. All Dravet animals demonstrated a marked hippocampal overexpression, particularly evident in CA1 stratum pyramidale neurons (Fig. 7A-B).

**Fig. 7.**
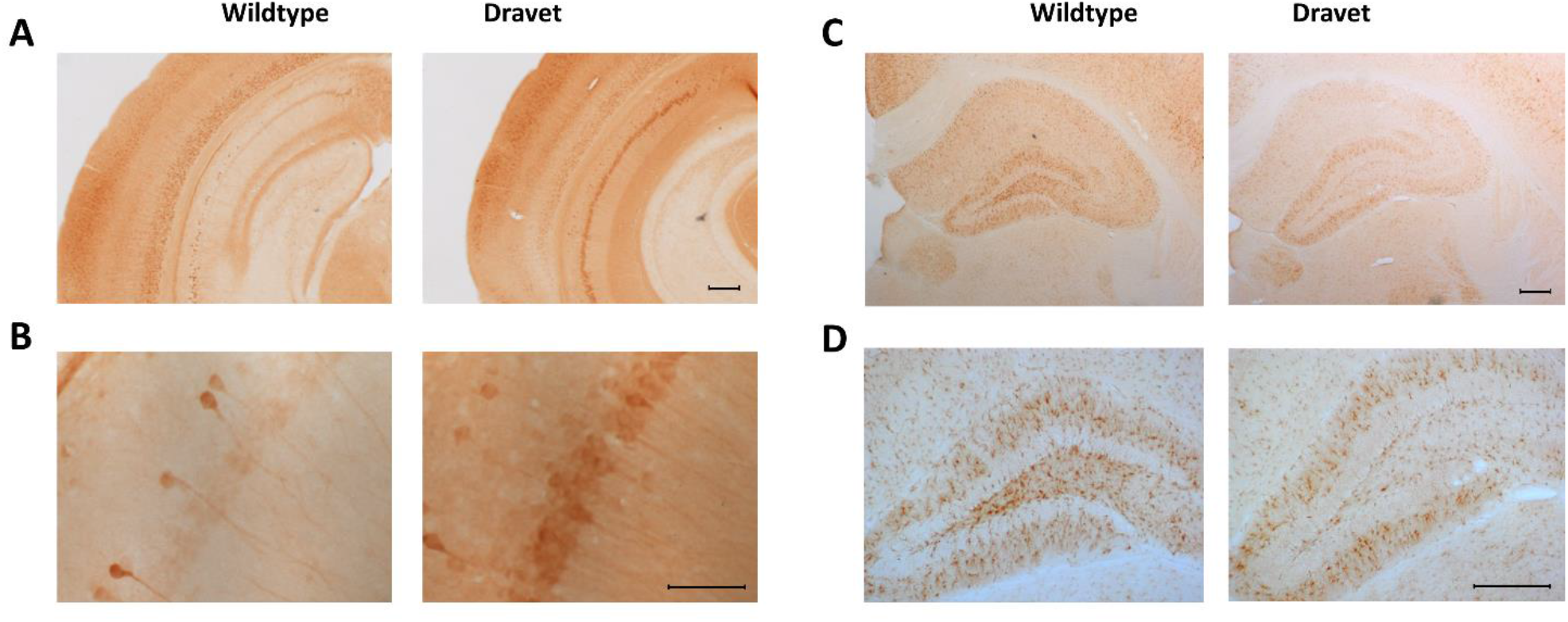
Immunohistochemical staining of representative proteins. **A-B** Representative immunohistochemical staining of PPP1R1B protein in the hippocampus (low magnification, **A**) and hippocampal CA1 region (high magnification, **B**). Pronounced protein immunostaining was particularly evident in the CA1 region of Dravet mice. **C-D** Immunohistochemical staining of HSD11B1 protein in the hippocampus (low magnification, **C**) and hippocampal hilar region (higher magnification, **D**). Protein immunoreactivity was reduced in Dravet mice, which was particularly evident in the hilus. Scale bars = 200 µm (A, C, D) and 50 µm (B).

In four-week-old mice, pathway enrichment analysis also demonstrated an over-representation of proteins functionally linked to calcium signaling (q=1.4e^-04^). Individual protein changes pointed to a down-regulation of calcium channel proteins and calcium transporters (Fig. 6B-k). Calcium/calmodulin-dependent protein kinases (Fig. 6B-l) were reduced in Dravet mice.

The list of regulated pathways also included a pathway involved in secondary cell signaling (*Phosphatidylinositol signaling pathway*, q=0.008). Moreover, an over-representation of proteins linked with the *nitric oxide (NO) signaling pathway* (q=0.005) was evident. In this context, the increased level of nitric oxide synthase, the main enzyme responsible for NO synthesis from L-arginine (Knowles et al., 1994), is of interest.

In addition to pathway enrichment analysis, we also identified the proteins with the most prominent regulation pattern. The strongest up-regulation was evident for the intermediate filament proteins glial fibrillary acidic protein (GFAP) and vimentin. The expression of both proteins was elevated at least two-fold in Dravet mice. The proteins with the strongest down-regulation were TRIM32 and HSD11B1. Proteomic profiling pointed to a reduction in HSD11B1 expression in Dravet mice, with a two times lower abundance than in wildtype mice. This finding was further confirmed by immunohistochemistry. An apparent down-regulation of the protein was evident in the hippocampus with the most obvious reduction in the hilus (Fig. 7C-D).

## Discussion

The large-scale proteomic analysis in a novel conditional mouse model of Dravet syndrome with *Scn1a* genetic deficiency provided comprehensive information about the molecular alterations characterizing different disease phases. Respective information about the proteomic signature of Dravet syndrome suggests possible pathophysiological mechanisms beyond *Scn1a* haploinsufficiency that may be involved in epileptogenesis and ictogenesis in Dravet mice.

As a basis for the proteomic analysis, we initially aimed to validate the novel conditional knock-in mouse model of Dravet syndrome with a heterozygous *Scn1a*-A1783V mutation. A model with this mutation has previously been generated on a pure C57BL/6J background (Ricobaraza et al., 2019) and a mixed (90:10) C57BL/6J and 129S1 background (Kuo et al., 2019) resulting in a more severe phenotype and higher mortality rate. Here, we characterized the model bred on a mixed (50:50) C57BL/6J and 129S1 background with an *Hprt* promoter mediated neuronal knock-in. We confirmed the development of spontaneous seizures and an increased susceptibility to hyperthermia-induced seizures and demonstrated a mortality rate of 40 %. Besides the seizure phenotype, we also observed behavioral alterations dominated by hyperactivity, which seem to reflect hyperactivity and attention deficits as common behavioral symptoms in patients with Dravet syndrome (Battaglia et al., 2016; Besag, 2004; Dravet, 2011).

When compared to other animal models of Dravet syndrome with a heterozygous *Scn1a* mutation, our model showed a similar age for onset of spontaneous seizures, increased susceptibility to thermally provoked seizures and notable hyperactivity. The SUDEP rate was relatively low compared to other models and occurred almost exclusively within a short time window (Table A.2). With the approaches used in this study, we failed to detect motor and social deficits in Dravet mice, which have been reported in selected mouse models (Table A.2). However, an improved performance on the accelerated rotarod test was also found in another mouse model (Ito et al., 2013), suggesting the test itself may not be appropriate to assess the Dravet-associated alterations in motor function and coordination. In this context, it is of interest that we confirmed alterations in gait in a follow-up study with a catwalk test and a detailed assessment of gait (Miljanovic et al., under revision).

To our knowledge, this is the first report of anhedonia-associated behavior in a Dravet mouse model indicating that *SCN1A* deficiency may have an impact on the affective state and may predispose to depression. However, we cannot exclude the possibility that anhedonia-associated behavior may be influenced by a taste preference related to *Scn1a* genetic deficiency.

Information about body weight development has not been provided for all Dravet mouse models. As such, for several models, it is unclear whether there was no delay or whether body weight development was not assessed and documented. The impact of the genetic deficiency on body weight development has only been reported in selected Dravet mouse models with heterozygous (Ricobaraza et al., 2019) and homozygous mutations (Martin et al., 2010; Ogiwara et al., 2007). In line with these reports, we observed a transient delay in body weight around weaning with animals catching up within five (females) or 20 (males) days following weaning.

### GABAergic, glutamatergic, and dopaminergic neurotransmission

Loss of function of sodium channel subunits encoded by the *SCN1A* gene in GABAergic interneurons is considered as the main source of hyperexcitability and ictogenesis in Dravet patients (Brunklaus and Zuberi, 2014; Catterall, 2018) and animals carrying the respective mutation (Almog et al., 2019; Kalume et al., 2015; Mantegazza and Broccoli, 2019; Mistry et al., 2014; Rubinstein et al., 2015; Salgueiro-Pereira et al., 2019; Tai et al., 2014; Tsai et al., 2015; Yu et al., 2006). However, an impact of cellular consequences in excitatory neurons on seizure susceptibility has also been suggested based on experimental findings. For instance, hyperexcitability of dissociated hippocampal pyramidal neurons (Mistry et al., 2014) and granule cells in the dentate gyrus (Tsai et al., 2015) in the period of chronic epilepsy may promote life-threatening seizures, known to exacerbate mice phenotype (Dutton et al., 2017; Salgueiro-Pereira et al., 2019). On the other hand, Ogiwara and colleagues demonstrated that *Scn1a* haploinsufficiency in hippocampal excitatory neurons can ameliorate seizures in Dravet mice (Ogiwara et al., 2013). Interestingly, Almog and colleagues showed how the hyperexcitability of CA1 pyramidal neurons during the pre-epileptic state can switch to hypoexcitability during the epileptic state in Dravet mice suggesting a role for these neurons in seizure propagation (Almog et al., 2019).

Altogether, these data suggest that both inhibitory and excitatory neurotransmission may be directly and indirectly affected by NaV1.1 dysfunction. Interestingly, the proteomic data suggest differential expression of multiple proteins linked with inhibitory and excitatory neurotransmission in *Scn1a*^+/-^ mice.

With changes in the abundance of various GABAA and GABAB receptor subunits, our findings indicate that signaling via both GABA receptor systems can be altered as a consequence of a *Scn1a* genetic deficiency with potential consequences for phasic and tonic inhibition. It should therefore be considered that both an up- and a down-regulation was observed for the different receptor subunits.

Concerning glutamatergic signaling, proteomic patterns in Dravet mice revealed a comprehensive down-regulation of subunits of NMDA, AMPA, and kainate glutamate receptors. In this context, it is of additional interest that several proteins linked with NMDA receptor function in the post-synaptic density showed a decreased abundance in Dravet mice. In addition, SYNGAP1, a post-synaptic density protein that negatively modulates trafficking of AMPA receptors to the membrane (Jeyabalan and Clement, 2016), also exhibited a dysregulation in the hippocampus as a consequence of the *Scn1a* deficiency.

Concerning the expression patterns of metabotropic glutamate receptor proteins, it should be considered that some of these receptors have a negative feedback function that limits excessive glutamate secretion (Dedeurwaerdere et al., 2015). Along this line, the reduced expression of the class two metabotropic glutamate receptor proteins mGluR2 and mGluR3 in Dravet mice might be of functional interest.

In addition to alterations in receptor proteins and PSD proteins, the lowered abundance of glutamine synthase and higher abundance of CTPS2, which contributes to deamination of glutamine to glutamate (Kassel et al., 2010), may imply that changes occur in glutamate metabolism.

Taken together our proteomic data set suggests that complex alterations occur affecting GABA and glutamatergic signaling. The direction of the alterations seems to suggest that some of these changes may contribute to hyperexcitability, whereas others may rather reflect compensatory mechanisms. Further research is necessary to explore the potential functional consequences.

Depending on the receptor subtype, dopaminergic signaling can affect seizure thresholds (Bozzi and Borrelli, 2013). Altered abundance of the dopamine metabolizing enzymes MAOA and MAOB, of a downstream effector protein of D1 receptors (Bozzi and Borrelli, 2013; O’Sullivan et al., 2008), and of proteins stabilizing D2 and D3 receptors, suggests that it may also be of interest to assess dopamine concentrations in the brain of Dravet mice.

### Voltage-gated ion channels

Voltage-gated ion channels affect neuronal excitability in different subcellular localizations therefore serving as important target sites for different antiseizure drugs (Sills and Rogawski, 2020). At the presynaptic level, P/Q-and N-type calcium channels represent important regulators of neurotransmitter release (Kassel et al., 2010). Thus, the extensive down-regulation of voltage-gated calcium channel subunits may constitute a compensatory mechanism counteracting the increased neuronal excitability characterizing Dravet syndrome. In this context, it is of additional interest that various calcium/calmodulin-dependent protein kinase (CaMK) subtypes proved to be reduced in hippocampal tissue from Dravet mice. CaMKs are important regulators that translate intracellular calcium concentrations into phosphorylation patterns with functional consequences for the targeted proteins (Swulius and Waxham, 2008).

Several potassium channels regulate outward potassium currents, thereby affecting membrane polarization and neuronal excitability (Villa and Combi, 2016). Proteomic data revealed a reduction of three potassium channel subunits (Kv1.2, Kv2.1 and Kv4.2) that attenuate back-propagating action potentials and prevent highly repetitive neuronal firing as one of the pathophysiological hallmarks of epileptic seizures (Niday and Tzingounis, 2018).

In summary, various changes in voltage-gated ion channel proteins occur in the Dravet mouse model. The reduction of calcium channel subunits and of potassium channel subunits may have contrasting consequences, which should be assessed in follow-up investigations.

While previous studies have also suggested alterations in sodium channel subunits with an up-regulation of NaV1.3 in hippocampal interneurons in a different Dravet mouse model (Yu et al., 2006), our data did not detect this protein in the *Scn1a*-A1783V mouse model.

### Astrogliosis, angiogenesis and NO signaling

Reactive astrogliosis can promote hyperexcitability, affect inflammatory signaling, and disrupt integrity of the blood–brain barrier (Devinsky et al., 2013). Increased GFAP abundance provides evidence for astrogliosis in the *Scn1a*-A1783V mouse model. This finding is in line with previous reports describing astrogliosis in other mouse models of Dravet syndrome (Alonso Gómez et al., 2018; Hawkins et al., 2019). Alterations in the astrocytic functional state are further supported by evidence of a reduction of astroglial excitatory amino acid transporters (EAAT1 and EAAT2) and for an overexpression of the inward rectifying potassium channel Kir4.1 (KCNJ10). These findings may imply contrasting functional consequences; however, this requires confirmation by further investigations.

Following seizure onset, we observed an additional regulation of ANXA2 and vimentin, which both act as modulators of VEGF signaling and angiogenesis (Dave and Bayless, 2014; Liu and Hajjar, 2016), previously discussed as a pro-epileptogenic factors and as a potential target for antiepileptogenesis (Morin-Brureau et al., 2012; Rigau et al., 2007).

Proteomic profiling also revealed a dysregulation of nitric oxide (NO) signaling in Dravet mice with an increased expression of neuronal nitric oxide synthase (nNOS) following the onset of spontaneous recurrent seizures. Considering that NO can induce reactive glial proliferation and promote angiogenesis (Arhan et al., 2011; Morbidelli et al., 2004), dysregulated NO signaling might play a role in epileptogenesis and promoting hyperexcitability in the epileptic brain.

### Proteomic alterations before epilepsy manifestation

While rather limited alterations were evident prior to epilepsy manifestation, they may be of interest as they might provide information about molecular mechanisms that occur as an early consequence of the *SCN1A* genetic deficiency thus potentially contributing to disease onset. In this context, the down-regulation of RASGRF1 might be of functional relevance as a contribution of RASGRF1 to epileptogenesis has been suggested based on a study with genetic and pharmacological targeting (Bao et al., 2018). Thus, it might be of interest to further explore a potential contribution of early RASGRF1 down-regulation to disease manifestation and seizure onset in Dravet mice.

### Differentially expressed proteins across time points

Proteome alterations in Dravet mice at the earlier time point were milder. Thereby the majority of dysregulated proteins were up-regulated. These molecular changes may represent a direct consequence of the *Scn1a* deficiency. In contrast, proteomic data from Dravet mice following disease manifestation indicated more pronounced molecular alterations with the majority of dysregulated proteins showing a down-regulation. As discussed above, some of these alterations may not only be related to *Scn1a* genetic deficiency but may also represent a mechanism compensating for repetitive seizure activity or other pathophysiological mechanisms developing throughout the course of disease. Notwithstanding the observed differences in regulation, the molecular function of regulated proteins at both time points showed the same qualitative pattern with the majority of regulated proteins being classified as enzymes and binding molecules. Furthermore, the functional annotation of differently expressed proteins revealed metabolite interconversion and protein modifying enzymes as the most common classification groups at both time points. These commonalities are particularly apparent with glutamatergic signaling, as one of highly dysregulated cellular processes following disease manifestation (Fig. 8). Changes in the expression of proteins with binding and catalytic activity, SHANK3 and subunits of protein kinases A and C (PRKAR1A, PRKCA), also suggest a possible dysregulation of glutamatergic synaptic transmission prior to spontaneous seizure onset.

**Fig. 8.**
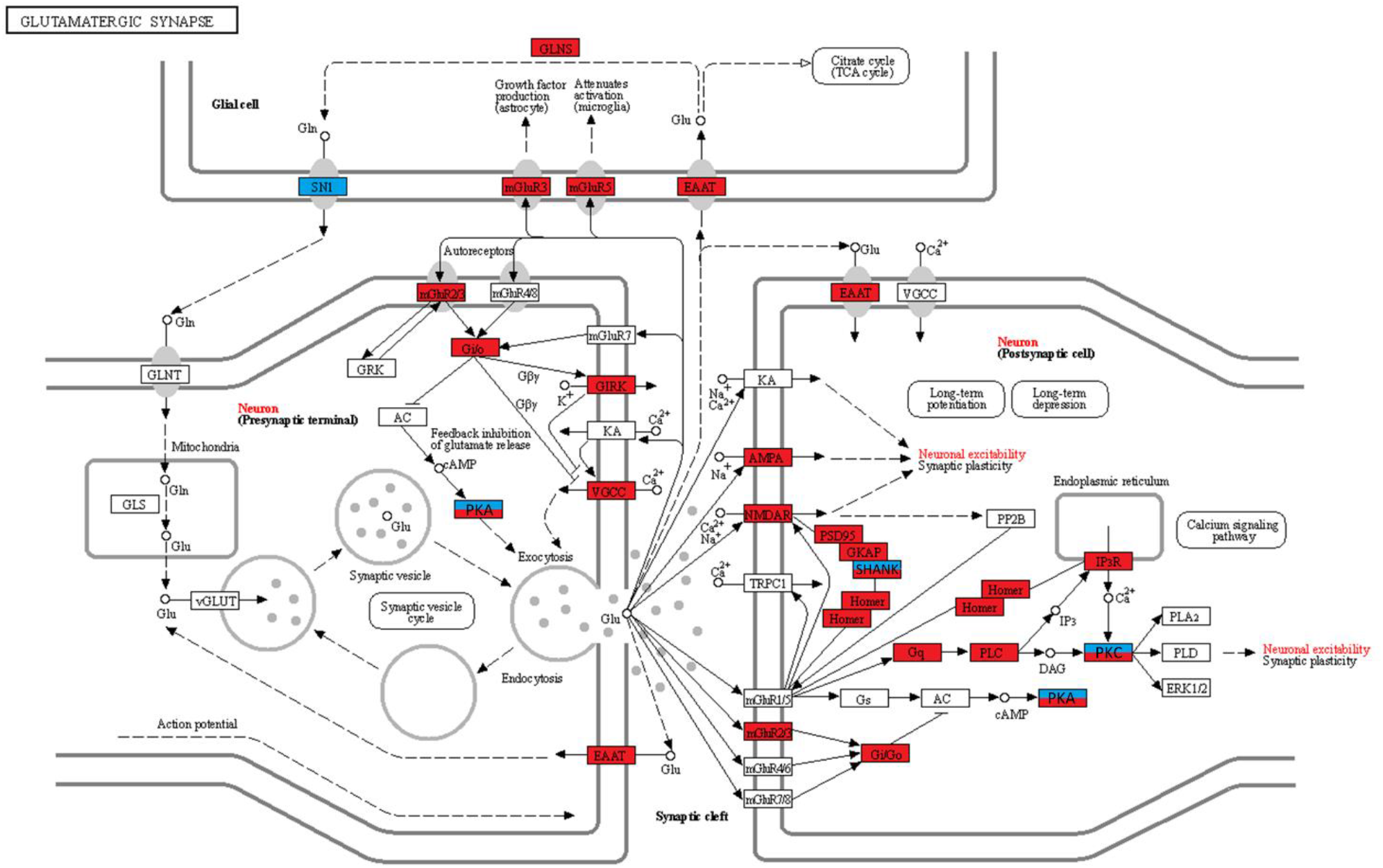
Glutamatergic synapse in Dravet mice (KEGG pathway). Significantly regulated proteins in two- and four-week-old Dravet mice are highlighted in blue and red, respectively. SHANK, PKA and PKC were the only proteins showing a dysregulation at the both time points, indicating that the dysregulation of glutamatergic synaptic transmission may being prior to disease manifestation. GLNS (GLUL) - glutamine synthetase; SN1 (SLC38A3) - solute carrier family 38 member 3; EAAT (EAAT1_2_3) – excitatory amino acid transporters 1, 2 and 3; mGluR - metabotropic glutamate receptor; GLNT (SLC38A1) - solute carrier family 38 member 1; GLS – glutaminase; vGLUT (SLC17A6_7_8) - solute carrier family 17 member 6, 7 and 8; GRK - beta-adrenergic-receptor kinase; Gi/Go (GNAI) - guanine nucleotide-binding protein G(i) subunit alpha; AC (ADCY1) – adenylate cyclase 1; PKA – protein kinase A; GIRK (KCNJ3) - potassium inwardly-rectifying channel subfamily J member 3; VGCC (CACNA1A) - voltage-dependent calcium channel P/Q type alpha-1A; KA (GRIK1) - glutamate ionotropic kainate receptor 1; AMPA (GRIA1) – glutamate receptor 1; NMDAR (GRIN1) - glutamate ionotropic receptor; TRPC1 - transient receptor potential cation channel subfamily C member 1; PSD95 – postsynaptic density protein 95; SHANK - SH3 and multiple ankyrin repeat domains protein; GKAP (DLGAP1) – discs large-associated protein 1; PP2B (PPP3C)- serine/threonine-protein phosphatase 2B catalytic subunit; Gq (GNAQ) - guanine nucleotide-binding protein G(q) subunit alpha; PLC (PLCB) - phosphatidylinositol phospholipase C beta; PKC (PRKCA) - classical protein kinase C alpha type; PLA2 (PLA2G4) - cytosolic phospholipase A2; IP3R (ITPR1) - inositol 1,4,5-triphosphate receptor type 1; PLD (PLD1_2) - phospholipase D1 and 2; ERK1/2 (MAPK1_3) - mitogen-activated protein kinase 1 and 3.

### Study limitations

Considering the whole proteome approach applied to hippocampal samples and the characteristics of the mouse model, respective limitations need to be considered.

Firstly, disease manifestation occurs early on in Dravet mice resulting in delayed postnatal bodyweight development. This in itself may impact molecular alterations in the brain.

Moreover, considering that untargeted proteomic studies are limited to screening proteins in the entire sample, further studies investigating the expression patterns in different hippocampal sub-regions and cell types along with studies addressing the functional consequences are needed to elucidate the functional relevance of these findings. The present findings provide a perfect basis to design respective studies applying targeted proteomic approaches. Finally, bulk approaches can also imply the risk to miss relevant changes in a selected cell population due to a dilution effect.

As a matter of course interpretation needs to take into account that molecular alterations can be a mere consequence of repeated seizure activity or disease-associated alterations without functional implications. Thus, as repeatedly highlighted, it is of utmost relevance to assess the functional implications of these findings in future studies.

## Conclusions

In conclusion, the whole proteome analysis in a mouse model of Dravet syndrome demonstrated complex molecular alterations in the hippocampus as a consequence of the genetic deficiency. Some of these alterations may have an impact on excitability or may serve compensatory functions, which should be further investigated.

Moreover, the proteomic data indicate that, due to the molecular consequences of the genetic deficiency, the pathophysiological mechanisms become more complex over the course of the disease. Resultantly, the management of Dravet syndrome may need to consider further molecular and cellular alterations. Functional follow-up studies are therefore required to confirm our findings and may provide valuable guidance for the development of novel therapeutic approaches.

## Acknowledgements

The authors thank Sarah Driebusch, Verena Buchecker, Helen Stirling, Fabio Wolf, Sieglinde Fischlein, Katharina Gabriel and Katharina Schönhoff for their excellent technical assistance.

We are very grateful to Ana Mingorance and the Dravet Syndrome Foundation in Spain for their efforts and investment to generate a commercially available Dravet mouse line in collaboration with the Jackson laboratory. Moreover, we thank Ana Mingorance for all her further support and the discussions about our planning and data.

This research is supported by a grant of the Deutsche Forschungsgemeinschaft (DFG, FOR 2591, GZ: PO681/9-1).

## Abbreviation list

MS: Mass spectrometry
LC-MS/MS: Liquid chromatography with tandem mass spectrometry
Hprt: Hypoxanthine-guanine phosphoribosyltransferase
VEGF: Vascular endothelial growth factor
JAK-STAT: The Janus kinase/signal transducers and activators of transcription
PSD: Postsynaptic density
CaMK: Calcium/calmodulin-dependent protein kinases n
NOS: Neuronal nitric oxide synthase
NO: Nitric oxide
SE: Status epilepticus

