## Appendix A. Supplementary Material for "Proteomic signature of the Dravet syndrome in the genetic *Scn1a*-A1783V mouse model"

**Table A.1. List of abbreviations (left column) and full names (right column) of all regulated proteins mentioned in the manuscript and their general function.**

| ***GABAergic signaling*** | |
| --- | --- |
| GABBR1 | Gamma-aminobutyric acid type B receptor subunit 1 |
| GABBR2 | Gamma-aminobutyric acid type B receptor subunit 2 |
| GABRA1 | Gamma-aminobutyric acid receptor subunit alpha-1 |
| GABRB1 | Gamma-aminobutyric acid receptor subunit beta-1 |
| GABRB3 | Gamma-aminobutyric acid receptor subunit beta-3 |
| SLC6A1 | Sodium- and chloride-dependent GABA transporter 3 |
| ***Glutamatergic signaling*** | |
| CTPS2 | Cytidine triphosphate synthetase 2 |
| GluA1 | Glutamate ionotropic receptor AMPA subunit 1 |
| GluA2 | Glutamate ionotropic receptor AMPA subunit 2 |
| GluA3 | Glutamate ionotropic receptor AMPA subunit 3 |
| GluA4 | Glutamate ionotropic receptor AMPA subunit 4 |
| GluK2 | Glutamate ionotropic receptor kainate type subunit 2 |
| GLUL | Glutamine synthetase |
| GluN1 | Glutamate ionotropic receptor NMDA type subunit 1 |
| GluN2A | Glutamate ionotropic receptor NMDA type subunit 2A |
| GluN2B | Glutamate ionotropic receptor NMDA type subunit 2B |
| mGluR2 | Metabotropic glutamate receptor type 2 |
| mGluR3 | Metabotropic glutamate receptor type 3 |
| mGluR5 | Metabotropic glutamate receptor type 5 |
| ***Synaptic transmission*** | |
| DLG2 | Disks large homolog 2 |
| DLG3 | Disks large homolog 3 |
| DLG4 | Disks large homolog 4 |
| DLGAP1 | Disks large-associated protein 1 |
| DLGAP2 | Disks large-associated protein 2 |
| DLGAP4 | Disks large-associated protein 4 |
| HOMER2 | Homer Scaffold Protein 2 |
| NLGN2 | Neuroligin 2 |
| NLGN3 | Neuroligin 3 |
| PCLO | Protein piccolo |
| PPP3CA | Serine/threonine-protein phosphatase 2B catalytic subunit alpha isoform |
| PPP3CB | Serine/threonine-protein phosphatase 2B catalytic subunit beta isoform |
| PRKCG | Protein kinase C gamma type |
| RASGRF1 | Ras-specific guanine nucleotide releasing factor 1 |
| RIMS1 | Regulating Synaptic Membrane Exocytosis 1 |
| SHANK1 | SH3 and multiple ankyrin repeat domains protein 1 |
| SHANK2 | SH3 and multiple ankyrin repeat domains protein 2 |
| SHANK3 | SH3 and multiple ankyrin repeat domains protein 3 |
| SYNGAP1 | Ras/Rap GTPase-activating protein SynGAP |
| SYT | Synaptotagmin |
| RPH3A | Rabphilin-3A |
| BAIAP2 | Brain-specific angiogenesis inhibitor 1-associated protein 2 |
| ***Dopaminergic signaling*** | |
| EPB41L1 | Band 4.1-like protein 1 |
| EPB41L3 | Band 4.1-like protein 3 |
| GNG7 | Guanine nucleotide-binding protein G(I)/G(S)/G(O) subunit gamma-7 |
| MAOA | Monoamine oxidase A |
| MAOB | Monoamine oxidase B |
| PPP1R1B | Protein phosphatase 1 regulatory subunit 1B |
| ***Calcium signaling*** | |
| CACNA1A | Calcium voltage-gated channel subunit alpha1 A |
| CACNA1E | Calcium voltage-gated channel subunit alpha1 E |
| CACNA2D3 | Calcium voltage-gated channel auxiliary subunit alpha2delta 3 |
| CACNB3 | Calcium voltage-gated channel auxiliary subunit beta 3 |
| CACNB4 | Calcium voltage-gated channel auxiliary subunit beta 4 |
| CACNG8 | Calcium voltage-gated channel auxiliary subunit gamma 8 |
| CAMK1D | Calcium/calmodulin-dependent protein kinase type 1D |
| CAMK2A | Calcium/calmodulin-dependent protein kinase type II subunit alpha |
| CAMK2B | Calcium/calmodulin-dependent protein kinase type II subunit beta |
| CAMK4 | Calcium/calmodulin-dependent protein kinase type IV |
| CAMKK1 | Calcium/calmodulin-dependent protein kinase kinase 1 |
| CAMKK2 | Calcium/calmodulin-dependent protein kinase kinase 2 |
| ***Potassium channels*** | |
| KCNA2 | Potassium voltage-gated channel subfamily A member 2 |
| KCNAB2 | Voltage-gated potassium channel subunit beta-2 |
| KCND2 | Potassium voltage-gated channel subfamily D member 2 |
| KCNJ10 | ATP-sensitive inward rectifier potassium channel 10 |
| KCNJ3 | G protein-activated inward rectifier potassium channel 1 |
| KCNJ9 | G protein-activated inward rectifier potassium channel 3 |
| ***Reactive astrogliosis*** | |
| GFAP | glial fibrillary acidic protein |
| EAAT1 | Excitatory amino acid transporter 1 |
| EAAT2 | Excitatory amino acid transporter 2 |
| ***Angiogenesis*** | |
| ANXA2 | Annexin A2 |
| ANXA4 | Annexin A4 |
| ITGAV | Integrin alpha-V |
| KDR | Vascular endothelial growth factor receptor 2 |
| VIM | Vimentin |
| ***Nitric oxide signaling*** | |
| NOS1 | Neuronal nitric oxide synthase |
| ***Cytoskeletal proteins*** | |
| ADD2 | Beta-adducin |
| ARPC1A | Actin-related protein 2/3 complex subunit 1A |
| CKAP4 | Cytoskeleton-associated protein 5 |
| CTTNBP2 | Cortactin-binding protein 2 |
| DBN1 | Drebrin |
| EZR | Ezrin |
| MAP1A | Microtubule-associated protein 1A |
| MYO6 | Unconventional myosin-VI |
| PLEC | Plectin |

**Table A.2. A comparison between here characterized mouse model of Dravet syndrome (*Scn1a*-A1873V, the first row) and other available mouse models with a heterozygous *Scn1a* mutation.** The arrows indicate the direction of observed phenotypic alterations. Minus (-) indicates no significant change in the evaluated parameter. A grey horizontal line was used between mouse models carrying the same *Scn1a* mutation, while a black horizontal line was used between mouse models carrying a different *Scn1a* mutation. P – postnatal day, PW – postnatal week, M – months, n.a. - not available, HIS – hyperthermia-induced seizures. ^1^(Ricobaraza et al., 2019); ^2^(Kuo et al., 2019); ^3^(Ogiwara et al., 2007); ^4^(Ito et al., 2013); ^5^(Dutton et al., 2013); ^6^(Han et al., 2012; Kalume, 2013; Oakley et al., 2009); ^7^(Yu et al., 2006); ^8^(Cheah et al., 2012); ^9^(Ogiwara et al., 2013); ^10^(Miller et al., 2014); ^11^(Mistry et al., 2014); ^12^(Tsai et al., 2015); ^13^(Dutton et al., 2017; Sawyer et al., 2016); ^14^(Martin et al., 2010).

| Mouse model | BL/6:129S1 background [~%] | Mortality rate [%] | Spontaneous seizures onset | HIS threshold, test day (P) | Activity level | Anxiety-like behavior | Social behavior | Cognition | Anhedonia-related behavior | Motor coordination | Body weight |
| --- | --- | --- | --- | --- | --- | --- | --- | --- | --- | --- | --- |
| *Scn1a*-A1783V | 50:50 | 40 | P16 | 39.4  (P23) | ↑ | ↓ | ↑ | - | ↑ | ↓ | ↓ |
| ^1^*Scn1a*^WT/A1783V^ | 100:0 | 75 | PW3 | 38.2  (1-6 M) | ↑ | ↑ | - | ↓ | n.a. | ↓ | ↓ |
| ^2^*Scn1a^∆E26^* | 90:10 | 100 | P14 | 41.1  (P12-14) | n.a | n.a. | n.a. | n.a. | n.a. | n.a. | - |
| ^3^ *Scn1a*^RX/+^ | 75:25 | 40 | P18 | n.a. | n.a. | n.a. | n.a. | n.a. | n.a. | - | - |
| *^4^Scn1a*^RX/+^ | 100:0 | 40 | P18 | n.a. | ↑ | ↓ | ↓ | ↓ | n.a. | - | - |
| ^5^*Scn1a*^Flox/+^Cre^+/-^ | 100:0 | 100 | P21 | 40.7  (P22) | ↑ | ↑ | ↓ | ↓ | n.a. | n.a. | n.a. |
| ^6^*Scn1a*^+/-^ | 99.9:0.1 | 40 | P21 | 39.5  (P20-46) | ↑ | ↑ | ↓ | ↓ | n.a. | ↓ | n.a. |
| ^7^*Scn1a*^+/-^ | 0:100 | 10 | P21 | n.a. | n.a. | n.a. | n.a. | n.a. | n.a. | n.a. | n.a. |
| ^7^*Scn1a*^+/-^ | 100:0 | 80 | P21 | n.a. | n.a. | n.a. | n.a. | n.a. | n.a. | n.a. | n.a. |
| ^8^Scn1a^fl/+^ | 100:0 | 70 | P18 | 39  (P35) | ↑ | n.a. | ↓ | ↓ | n.a. | n.a. | n.a. |
| ^9^*Scn1a*^d/+^ | 97:3 | 25 | PW3 | n.a. | n.a. | n.a. | n.a. | n.a. | n.a. | - | n.a. |
| ^10^*Scn1a^tm1Kea^* | 75:25 | 54 | P24 | n.a. | n.a. | n.a. | n.a. | n.a. | n.a. | n.a. | n.a. |
| ^11^*Scn1a^tm1Kea^* | 50:50 | 50 | P18 | n.a. | n.a. | n.a. | n.a. | n.a. | n.a. | n.a. | n.a. |
| ^12^Scn1a^E1099X/+^ | 75:25 | 46 | P20 | 40.2  (PW3-5) | n.a. | n.a. | n.a. | n.a. | n.a. | n.a. | n.a. |
| ^13^*Scn1a*^RH/+^ | 100:0 | 5 | n.a. | 41.3  (P14-15) | ↑ | - | ↓ | ↓ | n.a. | ↓ | - |
| ^14^*Scn1a*^RH/+^ | mix | 5 | n.a. | 43.1  (P14-15) | n.a. | n.a. | n.a. | n.a. | n.a. | n.a. | - |

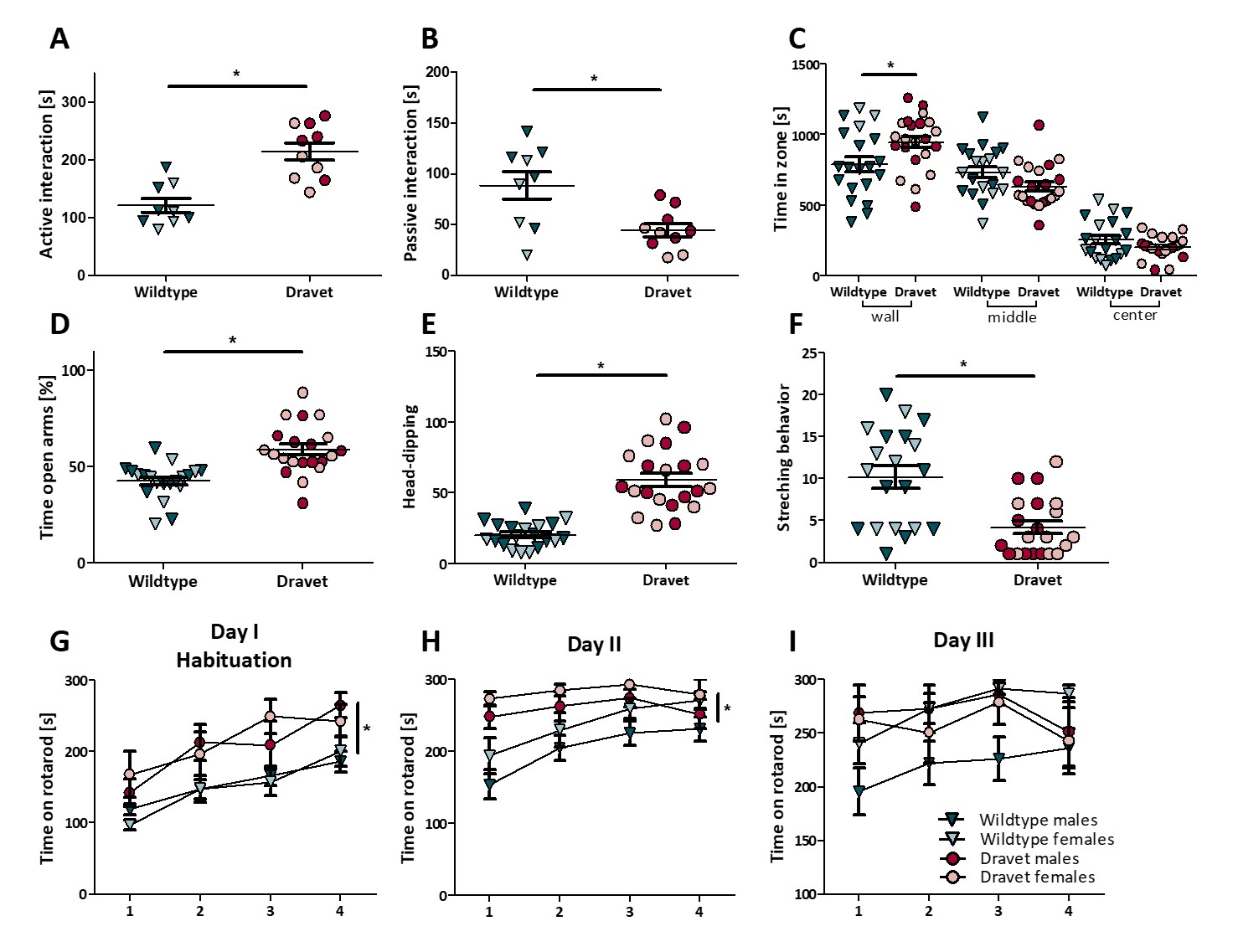
**Fig. A.1. Social interaction, thigmotaxic behavior, elevated plus maze and accelerated rotarod test. A** Time spent in active social interaction. Dravet mice engaged more in active interaction than their wildtype littermates. **B** Time spent in passive social interaction. Dravet mice spent less time than wildtype animals engaged in passive interaction. **A-B** Data shown are from 10 animal pairs with a Dravet genotype vs 9 wildtype animal pairs (Two-way ANOVA, Bonferroni post hoc; * p < 0.05, mean ± SEM). **C** Time spent in wall, middle and center zone over 30 minutes. Dravet mice spent more time in the wall zone than wildtype mice, while no significant difference was observed in the time spent in the middle or center zone (Three-way RM ANOVA, Bonferroni post hoc; * p < 0.05, mean ± SEM). **D** Time spent exploring open arms of elevated plus maze. Dravet mice exhibited a significantly higher preference towards open arms of the maze as compared to wildtype group. **E** Frequency to dip head over the maze. Dravet animals made significantly more head dips than wildtype mice. **F** Stretching postures to explore open arms of EPM. Dravet animals showed significantly lower number of stretching positions as compared to wildtype mice (Three-way ANOVA, Bonferroni post hoc; * p < 0.05, mean ± SEM). **G-I** Time on accelerated rod on three consecutive test days. Dravet mice performed better on rotarod comparing to wildtype mice. Females performed better than males (Five-way RM ANOVA, Bonferroni post hoc; * p < 0.05, mean ± SEM). **C-I** Data shown are from 21 (**D-F**) or 22 (**C, G-I**) animals with a Dravet genotype (n = 10 or 11 males, n = 11 females) vs 20 wildtype (n = 11 males, n = 9 females) animals.
